# Integrative analysis of mutated genes and mutational processes reveals seven colorectal cancer subtypes

**DOI:** 10.1101/2020.05.18.101022

**Authors:** Hamed Dashti, Abdollah Dehzangi, Masroor Bayati, James Breen, Nigel Lovell, Diako Ebrahimi, Hamid R. Rabiee, Hamid Alinejad-Rokny

**Author notes:** Equal contribution (Hamed Dashti, Abdollah Dehzangi).

## Abstract

Colorectal cancer (CRC) is one of the leading causes of cancer-related deaths in the world. It has been reported that ∼10%-15% of individuals with colorectal cancer experience a causative mutation in the known susceptibility genes, highlighting the importance of identifying mutations for early detection in high risk individuals. Through extensive sequencing projects such as the International Cancer Genome Consortium (ICGC), a large number of somatic point mutations have been identified that can be used to identify cancer-associated genes, as well as the signature of mutational processes defined by the tri-nucleotide sequence context (motif) of mutated sites. Mutation is the hallmark of cancer genome, and many studies have reported cancer subtyping based on the type of frequently mutated genes, or the proportion of mutational processes, however, none of these cancer subtyping methods consider these features simultaneously. This highlights the need for a better and more inclusive subtype classification approach to enable biomarker discovery and thus inform drug development for CRC. In this study, we developed a statistical pipeline based on a novel concept ‘gene-motif’, which merges mutated gene information with tri-nucleotide motif of mutated sites, to identify cancer subtypes, in this case CRCs. Our analysis identified for the first time, 3,131 gene-motif combinations that were significantly mutated in 536 ICGC colorectal cancer samples compared to other cancer types, identifying seven CRC subtypes with distinguishable phenotypes and biomarkers. Interestingly, we identified several genes that were mutated in multiple subtypes but with unique sequence contexts. Taken together, our results highlight the importance of considering both the mutation type and mutated genes in identification of cancer subtypes and cancer biomarkers.

## Introduction

Cancer is the leading cause of death worldwide, with colorectal cancer ranking second after lung cancer in lethality [1]. The Cancer Genome Atlas (TCGA) reported that 16% of CRC samples show DNA damage mechanisms and high tumor mutational burden [1]. Accurate identification and quantification of CRC subtypes, and delineating underlying molecular mechanisms, is an essential first step to help develop personalized medicine strategies leading to effective treatment modalities and better clinical outcomes for patients. There is currently no consensus on the number of CRC subtypes, and different approaches used to classify CRCs based on mutation or gene expression have reported up to six subtypes (refs). In 2013, Felipe De Sousa *et al.* [2] used a hierarchical clustering method on gene expression data from 90 CRC patients to identify three CRC subtypes (CCS1, CCS2, and CCS3). In the same year, Sadanandam *et al.* [3] analyzed gene expression data from a larger cohort comprising 445 patients, identifying five CRC subtypes [3].

In another study on gene expression data from 566 patients, Marisa *et al.* applied hierarchical clustering using the Ward linkage method [4] to identify CRC subtypes. By adopting the Pearson correlation coefficient (PCC) as a distance measure on the gene expression profiles they identified six CRC subtypes [4].

In another study on gene expression data from a cohort of 188 CRC patients, Roepman *et al*. identified three subtypes using an unsupervised clustering (hierarchical clustering) method [5]. Most recently, Guinney *et al*. [6] conducted a comprehensive study using gene expression data to identify CRC subtypes as a part of the International Consortium to Generate Colorectal Cancer Subtypes (CRCSC). This study identified four biologically distinct CRC subtypes called CMSs (consensus molecular subtypes [7]), each having unique clinical and molecular markers. For example, they defined CMS1 as a hyper-mutations case and CMS2 has a high copy number alteration. This study covered 87% of all CRC cases as well as the other remaining uncharacterized samples. They integrated all the data from previous studies and used a random forest (RF) algorithm (using 5,972 genes) and fitted a decision tree to each bootstrap. As a result, their final classifier was built as an ensemble of each of these decision trees.

There are also some studies that used somatic point mutations and pathways instead of gene expression for cancer subtype identification. Somatic mutation is the hallmark of cancer and different mutational processes and genes involved are typically linked to distinct mechanisms of tumor development, from which subtypes could be identified. In addition, the stable nature of somatic mutations makes them good candidates for cancer subtype classification. Kuijjer *et al*. used mutation data sets of 6,406 samples from 23 cancer types from the Cancer Genome Atlas (TCGA). They hypothesized that mutation processes in cancer can be identical in different cancer types, i.e. they were not always tissue-specific. Their analysis identified nine subtypes across all cancers [8].

Somatic point mutations in cancers are caused through several mutational processes. In 2015, Alexandrov *et al*. [9] showed that there are 30 mutational signatures in cancers, most of them associated with a specific molecular mechanism to uncover the causality behind somatic point mutations across the genome. The proposed concept provided the importance of motif context in the analysis of somatic point mutations in cancer genomics. It is also known that transcription factors bind to their specific sequence motifs, therefore sequence context of cancer mutations can potentially change the expression and regularization of genes including cancer driver genes. Hence, quantifying somatic mutations with respect to their gene-motifs instead of genes can provide greater discriminatory power for identification of cancer subtypes. To the best of our knowledge, the context of mutations in highly mutated genes has not been used for cancer subtype classification and therapeutic biomarker identification. Based on the merits associated with somatic mutations, in this study we perform an integrative analysis using a “gene-motifs” concept to accurately identify CRC subtypes. An overview of the pipeline used in this study can be found in **Figure 1**. We hypothesize that subtypes can be determined by not only mutated genes but also by the mutated sequence motifs within the genes, thus called “gene-motifs”. Us candidate gene-motifs as features of the model, we filter out irrelevant or less informative features using a statistical model (fitting best beta-binomial distribution) to avoid bias and reduce noise in our data, and then use a model-based clustering approach [10] to classify our CRC samples into subtypes. Our analyses identify seven unique subtypes that show distinct links to symptoms and genotypes in CRC. We also identify novel colorectal-associated genes and specific mutated motifs, which represent different CRC subtypes. We also clearly demonstrate how some of the subtypes are region-based, supporting the study by Kan *et al.* [11].

**Figure 1.**
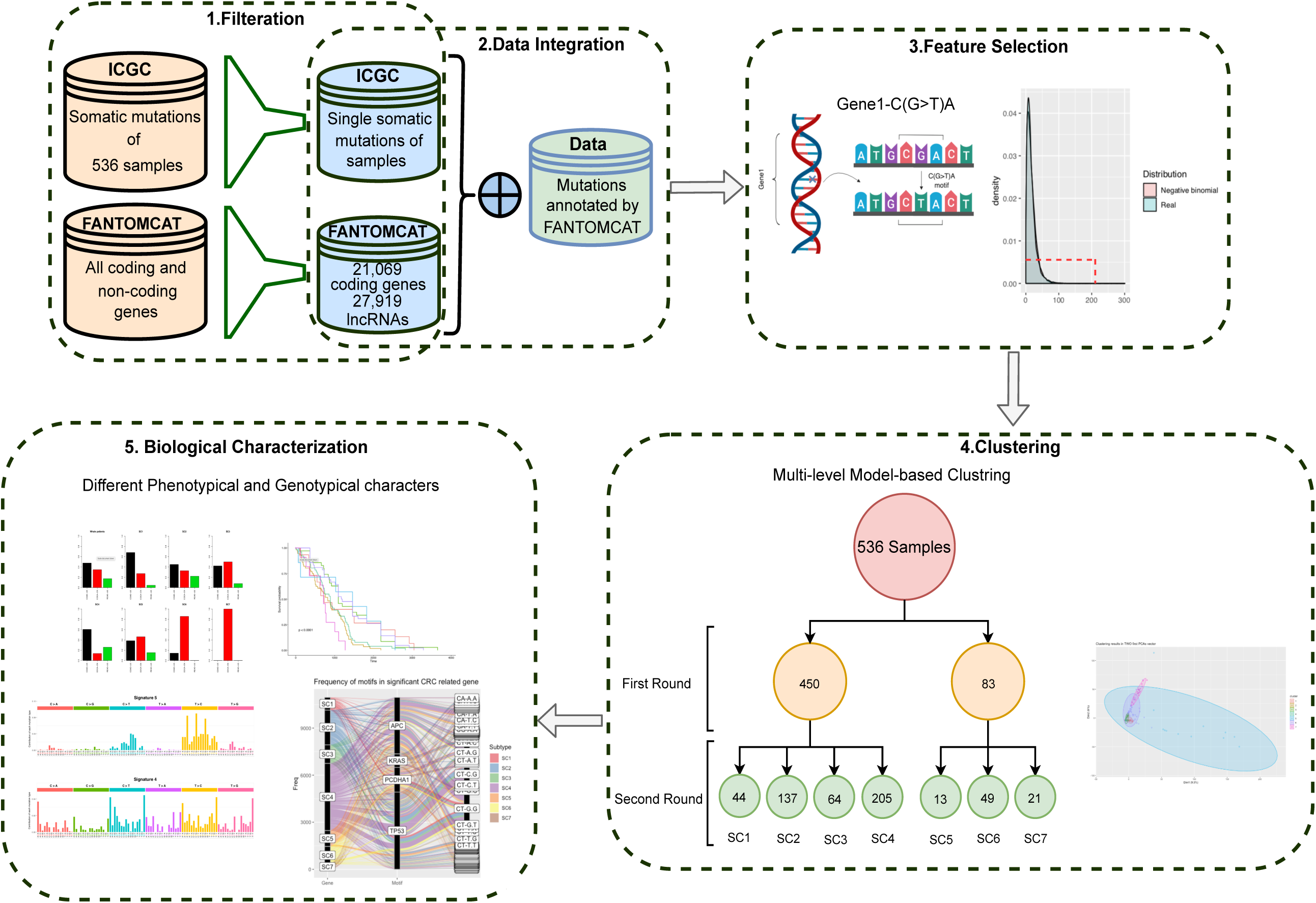
An overview of the pipeline used in this study. We downloaded publicly available somatic point mutations of 536 whole genome sequencing CRC individuals from the ICGC. We used the FANTOMCAT genes list to identify the number of mutations in coding and lncRNA genes. We then used negative-binomial and beta-binomial distributions to identify significantly mutated genes and gene-motifs, respectively. Using 3,131 candidate gene-motifs as the features of our model-based clustering, we identified seven subtypes in CRC individuals. Our comprehensive biological characterization showed that the identified subtypes have different mutational load in genes. Our mutational signature analysis also showed that different combinations of signatures become dominant in each subtype. We then identified genes and conserved motifs that significantly mutated in each subtype. Lastly, we performed gene ontology, pathway and survival curve analyses.

In addition to protein coding genes, non-coding RNA genes have been also demonstrated to act as key regulators of physiological diseases; particularly, long non-coding RNAs (lncRNAs) have been shown to act as oncogenic drivers and tumor suppressors in multiple cancers. In this study. we also investigate the difference in mutation rates between genes and lncRNAs, which can provide important insight into their overall impact on different CRC subtypes.

## Results

To build our unique clustering framework, we used somatic point mutations from the International Cancer Genome Consortium (ICGC), which contains data from 536 patients (45% female, 55% male) with CRC across three projects (READ-US, COAD-US, COCA-CN). Typically somatic point mutations are annotated within gene-coding regions exclusively, so we also extended annotations of somatic point mutations to 27,919 lncRNAs and 21,069 protein-coding genes annotated in the FANTOM CAGE associated transcriptome (FANTOM CAT) dataset [7].

### Statistical workflow to identify significant genes and gene-motifs

CRC samples have different mutational frequencies with diverse mutational profiles (**Supplementary figure S1**). To cluster these samples, we first identified candidate genes and gene-motifs (as the feature for our clustering) that were mutated at a significantly higher level in CRC samples relative to other cancers. For this purpose, we used a Cullen and Frey graph [12] with 100-fold bootstrapping to identify best-fitted distribution to our candidate genes data. Among different distributions, the negative-binomial distribution had the best-fitted distribution (**Supplementary figure S2**), which was validated by the Cramer-Von Mises test (*see the method section*). Using a negative-binomial distribution as the most suitable distribution to calculate the P-value of mutational load for each gene, we identified 382 significant protein coding genes that were significantly mutated (P-value < 0.01) in the CRC samples. The rankings of the significant coding genes based on their P-value (i.e. candidate genes) are provided in **Supplementary table S1**.

### Gene-motif concept

Having significantly mutated genes identified, we then tested different models to identify significantly mutated gene-motifs. the Gene-motifs refer to the sequence motifs mutated within a gene. Here, we define motif as a 3-nucleotide sequence context of the mutated nucleotide, i.e. NXN-to-NYN (where X has mutated to Y, and N: A, C, G, or T). There is in total 96 different mutations within all possible motifs in DNA [9]. We determine the number of each of these 96 mutations types within each gene. As described above, we investigated different models for fitting mutational profiles in gene-motifs and based on the comparison of Poisson, negative-binomial, Weibull, Gamma, log-normal and beta-binomial distributions in a Cullen and Frey Graph (**Supplementary figure S3**), we found that the beta-binomial distribution to be the most suitable distribution to identify significantly mutated motifs within the significant genes (e.g. gene-motif). As a result, we identified 3,131 gene-motifs with P-values smaller than the threshold (P-value <0.01) as significant gene-motifs, which have been used as the feature to cluster CRC patients. The list of candidate gene-motifs is provided in **Supplementary table S2**. An overview of the gene-motif concept is provided in **Supplementary figure S4.**

### Distinguishing subtypes based on the candidate gene-motifs

We used principal component analysis (PCA) to investigate the effectiveness and validity of our selected features. As it shown in **Supplementary figure S5**, the similar length for the first two principal components (PCs) demonstrates significant potential discriminatory information (the variance of data along these features is almost the same) that can be obtained using this distribution, indicating the importance of our selected features.

We then used multi-level model-based clustering [10] to identify subtypes in CRC samples. We used this technique over previous clustering methods such as hierarchical clustering due to its ability to build a formal model without relying on the need for random initialization. Traditional clustering models, such as a k-means algorithm, are randomly initialized while others like hierarchical clustering are heavily dependent on heuristics to build their model. Additionally, clustering models such as dbscan [13], hdbscan [14], and local shrinking-based clustering [15] require the user to specify the optimal number of clusters or other parameters. Model-based clustering used here does not have such requirements, enabling us to find the most suitable number of clusters based on the nature of the available data. To build this model we first applied clustering to all patients to produce the first level of clusters and then applied the model-based clustering on each cluster separately to detect if each cluster can in turn be divided into new meaningful groups (i.e. clusters with considerable number of patients; 1% of the total number of samples). We repeated this process until no more meaningful clusters were generated (see more details in the method section).

Using the described approach, we identified two clusters with 450 and 83 patients at the first clustering level. These two clusters were then divided into smaller, but still distinguishable, sub-groups in the second clustering level. At this level, the first cluster was divided into four sub-clusters with 44, 137, 64 and 205 number of patients while the second cluster was divided into three clusters with 13, 49 and 21 number of patients. This pattern did not continue to the third level of clustering. At the third level, most of our clusters did not break down into smaller clusters; in two cases (137-sample and 205-sample clusters) the clusters divided into a large cluster and few outliers. These outliers were close to the larger cluster and far from the rest of the clusters. In these cases, instead of introducing a new cluster, we investigated those samples as outliers to the same cluster. As a result, our iterative clustering method identifies seven subtypes namely, SC1, SC2, SC3, SC4, SC5, SC6 and SC7 for CRC based on somatic point mutations that were determined on 3,131 candidate gene-motifs. The patients in each cluster are shown in two dimensional PCAs of features in **Supplementary figure S6** and listed in **Supplementary table S3.** Interestingly, a clustered heatmap of the fraction of mutated gene-motifs in each subtype shows that most of the gene-motifs are subtype specific (**Figure 2**). The frequency of 3,131 gene-motifs in each subtype is provided in **Supplementary table S4.** For the rest of this paper, we will comprehensively investigate biological interpretability of our identified subtypes.

**Figure 2.**
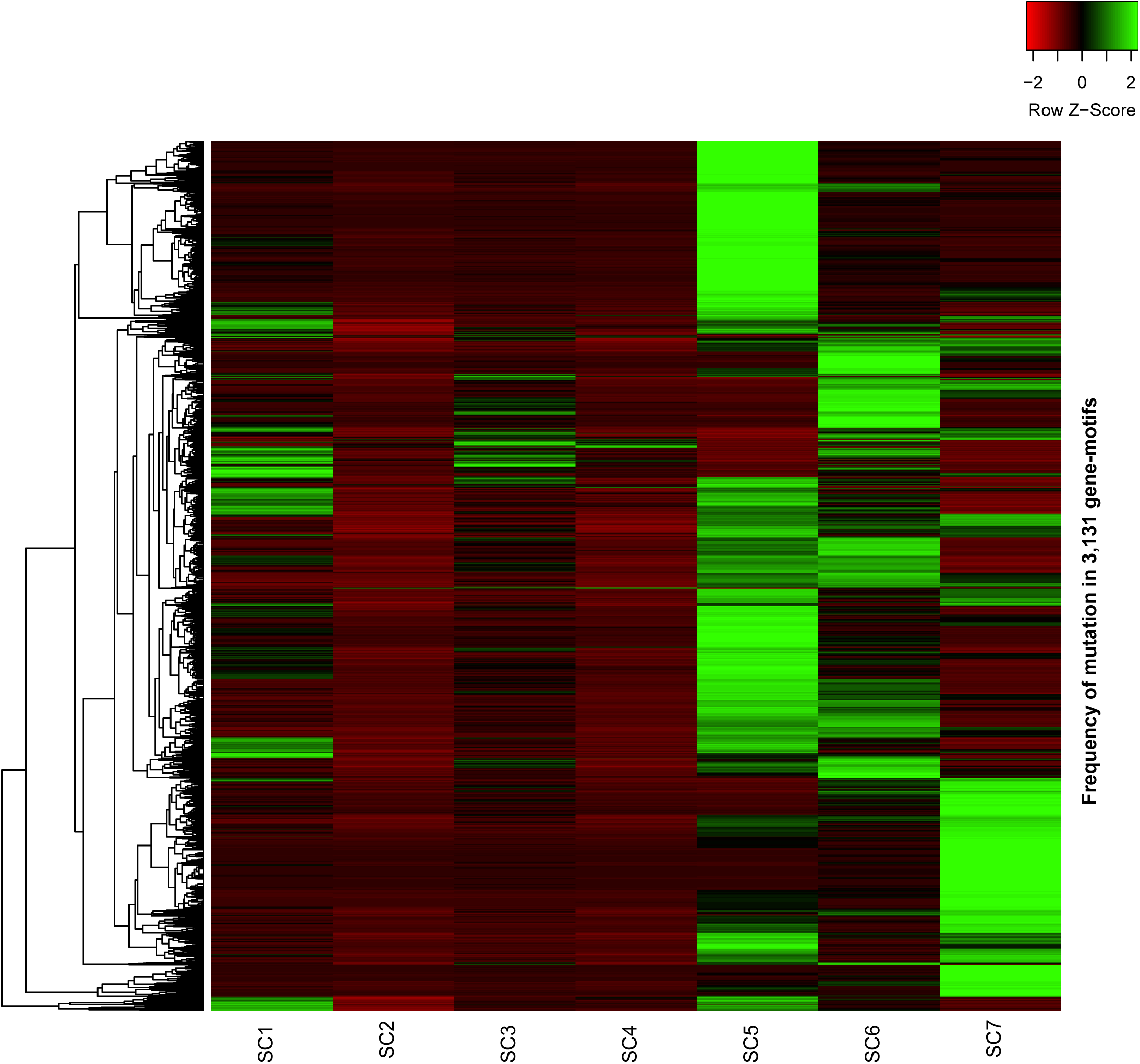
Clustered heatmap of 3,131 gene-motifs. Hierarchical clustering of 3,131 gene-motifs in seven CRC subtypes based on their mutational frequency. Intensity represents the frequency of mutation in each subtype.

### Biological characterization of each subtype

In this section, we comprehensively investigate the biological characterization of each CRC subtype to identify subtypes-specific genes, specific conserved motifs, and gene pathways which underlie colorectal cancer. We also mine the clinical data available for CRC patients.

### Gene association as the biomarker of each subtype

After determining each subtype, we comprehensively investigated the biological interpretability of each subtype by identifying subtype-specific genes. The 382 candidate genes revealed that many of them have been previously associated to CRC and/or other cancers. For example, we identified *PCDHGA6, TTN, TP53, FRG1BP, AP2A2, MUC6, H3BNH8, DOCK3, CACNA1D, SERPINB, ZBED6, ZBED6, NXPH1, NKAIN2*, and *EIF3H* genes that are known to be associated with CRC cancer [16-21]. We also used Fisher’s exact test to identify subtype-specific genes and also those genes that significantly mutated in multiple subtypes. We identified many common and subtype-specific genes, some of which are known to be associated with CRC or other cancers (**Supplementary table S5**). Interestingly, this analysis enables us to identify unique mutated genes in a specific subtype, in which none of the other subtypes have mutation in these genes (**Supplementary table S5**).

For example, *PCDHGA6* and *TTN* appeared as two of the significant genes in SC1. *PCDHGA6* was mutated in 42 of 44 patients of SC1 samples (95%), while mutated in 144 patients of 489 patients (29%), in other clusters. Most of the *PCDHG* family (e.g. *PCDHGB3* and *PCDHGA6*) are calcium-dependent cell-adhesion proteins [22], significantly mutated in SC1 patients and previously reported as diagnostic markers for cervical cancer [23]. *TTN*, which produces the very large protein called titin that has several functions within sarcomeres, was significantly mutated in above 90% of patients in SC1. One of the main function of this gene is to provide structure, flexibility, and stability of cell structures, which is associated to cancer [24]. We also found out that 90% of SC1 patients have been mutated in *CATG00000081433*, which is a FANTOMCAT-specific coding gene. Importantly, this gene encompasses multiple colorectal cancer associated GWAS (Genome-wide association study) SNPs. This gene is also dynamically expressed in an induced pluripotent stem cell model of neuron differentiation dataset [7].

*TP53* is the most mutated gene in SC2, which has been previously associated with a variety of cancers, including CRC [17]. Adenomatous polyposis coli (*APC)* was also found within this sub-cluster, a known driver gene for colorectal, pancreatic, desmoid, hepatoblastoma, and glioma cancers [25]. *APC* is a tumor suppressor gene that encodes a multi-functional protein that is involved in several processes, including regulation of β-catenin/Wnt signalling, intercellular cell adhesion, proliferation, apoptosis and differentiation [26].

In SC3, *FRG1BP*, a gene previously associated with prostate cancer [27], was mutated in ∼50% of samples in SC3 compared to less than 15% of samples in other clusters. *AP2A2* is another important gene that mutated in 54% of SC3 samples and in less than 23% of samples who are in other subtypes. *AP2A2*, as a component of the adaptor protein complex 2 (AP-2), is involved in clathrin-dependent endocytosis in which proteins are incorporated into vesicles surrounded by clathrin (clathrin-coated vesicles) [28]. More recently, a role for *AP2A2* has been suggested in asymmetric cell division and self-renewal of hematopoietic stem and progenitor cell [29]. Another important gene is *MUC6* which has mutated in 54% of SC3 samples and less than 24% in other samples. *MUC6* is shown to be associated with pancreatic ductal carcinoma and small intestine cancer [30]. It is also involved in “Defective *GALNT12* causes colorectal cancer 1 (*CRCS1*)” and “Defective *GALNT3* causes familial hyperphosphatemic tumoral calcinosis (*HFTC*)” pathways based on the Reactome Pathways database [31].

*H3BNH8* is the most mutated gene in SC4 samples (mutated in 61% of SC4 samples). This gene is an important paralog of APC (homologous genes that have diverged within one species). *TP53* and *APC* are other associated genes to this subtype. These genes are also significantly mutated in SC2 samples.

*DOCK3* has been mutated in 100% of SC5 samples but it is mutated in only 5% of samples in other subtypes. *DOCK3* is known as a modifier of cell adhesion (MOCA) and presenilin-binding protein (PBP) [32, 33], which has an important role in melanoma [34]. *CACNA1D* is another cancer related gene which is Calcium voltage-gated channel subunit alpha1 D [35] that is identified as an associated gene in samples of SC5. *SERPINB4* is another important cancer-related gene, which is mutated in 53% of SC5 samples and no mutation in other subtypes. The protein of *SERPINB4* can inactivate granzyme M, an enzyme that kills tumor cells [36]. *SERPINB4* has been previously identified as associated gene with loci 18q deletion syndrome and squamous cell carcinoma [37]. *ZBED6* is another important gene which has been mutated in 63% of SC5 and almost no mutation in other subtypes. *ZBED6* is known as a repressor of *IGF2* that influences cell proliferation and development that affects cell cycle and growth of human CRC cells [38].

*ZNF717* as one of the important genes that mutated in 85% of SC6 samples and only in 10% of samples in other subtypes. This gene involves many cell activities such as cell proliferation, differentiation and apoptosis, and in regulating viral replication and transcription [39]. *ZNF717* has been recently suggested as a candidate gene in the African-American population with CRC [39]. Interestingly, 90% of SC6 samples mutated in genes that are related to immunoglobulins such as the *IGHM* and *IGHV3* families.

Finally, we found *NXPH1* is mutated in all SC7 samples and mutated only in 2% of samples in other subtypes. This protein forms a very tight complex with alpha neurexins, a group of proteins that promote adhesion between dendrites and axons [40]. In addition, we identified 100% of SC7 samples also mutated in protein coding gene *CATG00000086870*, which was recently discovered by the FANTOMCAT consortia. Interestingly, this gene mutated in less than 1% of samples, who are in other subtypes. Moreover, using FANTOM5 expression atlas [41] we found this gene is highly expressed for colorectal tissue. *NKAIN2* is known as a tumor suppressor in Chinese prostate cancer [42]. This gene also mutated in all 21 Chinese samples of SC7 but only 2% of samples in other subtypes. *EIF3H* as a CRC potential diagnostic biomarker [43] is mutated in all SC7 samples but in only 3% of samples in other subtypes. *EIF3H* is a cancer-related gene and suggested as a CRC potential diagnostic biomarker [43]. We provided a ranked list (based on significant P-value) of the candidate genes in each subtype in **Supplementary table S5**.

### Mutational signature

Previous studies from Alexandrov *et al*. in 2015 identified that mutational signatures in cancer genomes demonstrate different biological mechanisms [9]. They identified 30 distinct mutational signatures of all cancers (mostly from whole exome sequencing) in which four (i.e. 1, 5, 6 and 10) are associated with CRC [9]. These CRC mutational signatures are associated to age of patients (signature 1), defective DNA mismatch repair (signature 6), altered activity of the error-prone polymerase POLE (signature 10); and the etiology of signature 5 is unknown [44]. To find which mutational processes are active in the CRC subtypes, we used the CANCERSIGN tool [45] to identify mutational signatures (*see the method section*). Using whole genome sequencing data from ICGC CRC samples, we identify seven signatures overall for the CRC patients (**Figure 3a**).

**Figure 3.**
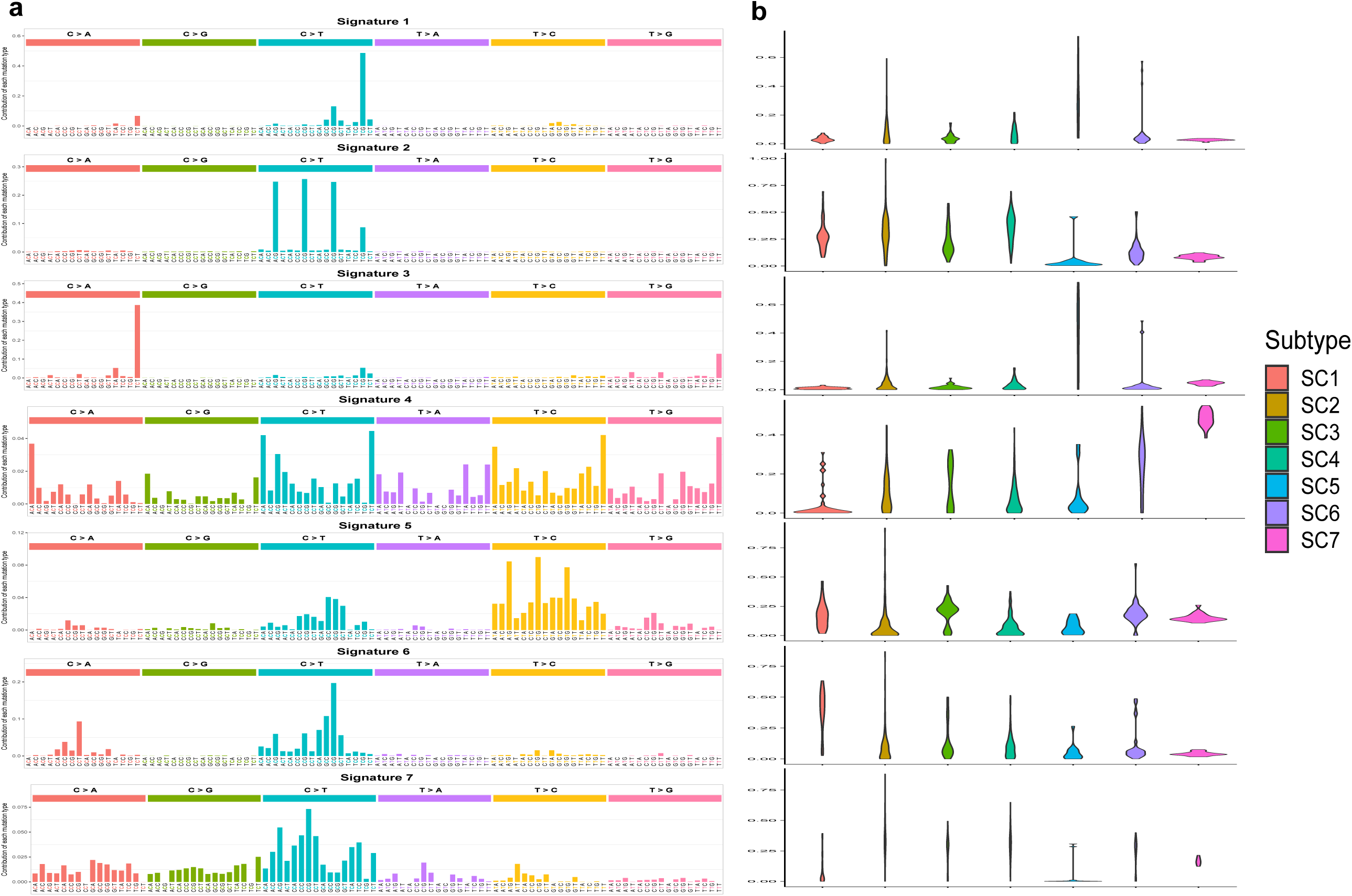
Cancer mutational signature analysis. We used the CANCERSIGN tool [58] to identify mutational signatures in each subtype. Using CANCERSIGN and WGS somatic mutation form the CRC samples, we identified seven mutational signatures where were very similar with those identified by Alexandrov *et al*. [9] except the signatures 1 and 3 that the linear combination of them can represent the signature 10 COSMIC (Alexandrov *et al.*). The exposure matrix for identified signatures are also provided in **Supplementary table S11**. The evaluation plot of deciphering 3-mer mutational signatures also provided in **Supplementary figure S11**.

Using the Pearson correlation, we calculated the correlation between identified signatures in our study and Alexandrov signatures (in **Supplementary figure S7**). As the figure shows there are strong correlations between the Alexandrov signatures and those signatures identified in this study. A linear combination of signatures 1 and 3 in our study (**Figure 3a**) represented COSMIC signature 10 which is also associated with CRC. Signature 2 in our study was a common signature between all subtypes (**Figure 2b**) and Signature 4 in our study is similar to signature 5 of Alexandrov (associated to CRC but no known etiology), but has a greater exposure in SC5. Signature 5 in our study is similar to signatures 16 and 26 Alexandrov which are not associated to CRC. The etiology of signature 16 is unknown, but signature 26 is possibly associated with defective DNA mismatch repair [44]. This signature has greater exposure in SC4 and SC5. Signature 6 in our study is similar to signature 6 Alexandrov, which is associated with CRC. This signature has greater exposure in SC3 and SC7 and possibly is associated with defective DNA mismatch repair [44]. Finally, signature 7 in our study is similar to signature 19 Alexandrov and is associated with pilocytic astrocytoma. The exposure of signature 7 is greater in SC2, SC3 and SC6 (see **Figure 3b** for more details). A detailed investigation of mutational load in coding and non-coding genes is also presented and discussed in the next section.

### Mutational rates of protein coding and lncRNA genes

We then investigate the mutation rate in protein-coding genes and lncRNAs of the clusters, which can provide important insight into their overall impact on different CRC subtypes. This can help us understand if our identified subtypes are different with respect to their mutational load in coding and non-coding genes. As is shown in **Supplementary figures S8** and **S9**, the mutation rate in coding and non-coding genes are different between subtypes. Interestingly, our analysis shows that SC5 is a hyper-mutated subtype (**Supplementary figure S8**). In addition, as it is shown in **Supplementary figure S10**, the mutational rate of long non-coding genes in subtype SC7 are much higher than other subtypes, suggesting a potential role for long non-coding RNA genes in this subtype.

We also investigated the difference between our identified subtypes with respect to the mutational load in different functional consequence or variant types (*see the method section*). Interestingly, we observed that mutations in some of the subtypes are enriched in different consequence types (**Supplementary figure S11**). For example, missense variants had a greater fraction in SC1, SC2, SC3 and SC4, while intron variants had a greater fraction in subtypes SC5, SC6 and SC7 (**Supplementary figure S11**). Interestingly, there is almost no missense variant in SC7, and our analysis also showing that most of the mutations in SC7 were intron variant (57.7%) and intergenic region variant (21.6%). Moreover, unlike other subtypes, there is no synonymous variant in SC7. In SC6, most of the mutations are in intron and downstream gene. A summary of this analysis is shown in **Table 1**.

**Table 1.** Consequence types of mutations in the CRC subtypes.

| Consequence | SC1 | SC2 | SC3 | SC4 | SC5 | SC6 | SC7 |
| --- | --- | --- | --- | --- | --- | --- | --- |
| 3_prime_UTR_variant | 3.08 | 3.21 | 2.5 | 2.94 | 2.94 | 2.28 | 0.38 |
| 5_prime_UTR_premature_start_codon_gain_variant | 0.19 | 0.16 | 0.18 | 0.23 | 0.16 | 0.14 | 0.02 |
| 5_prime_UTR_variant | 0.66 | 0.7 | 0.82 | 0.69 | 0.77 | 0.64 | 0.11 |
| downstream_gene_variant | 19.94 | 17.54 | 18.68 | 17.79 | 15.17 | 17.22 | 9.26 |
| exon_variant | 10.81 | 10.69 | 9.68 | 10.82 | 8.66 | 8.03 | 0.91 |
| initiator_codon_variant | 0 | 0.01 | 0 | 0 | 0 | 0 | 0 |
| intergenic_region | 0.18 | 0.06 | 1.45 | 0.15 | 2.67 | 5.56 | 21.58 |
| intragenic_variant | 0 | 0 | 0 | 0 | 0 | 0 | 0 |
| intron_variant | 14.65 | 15.46 | 24.61 | 12.63 | 27.78 | 31.56 | 57.41 |
| missense_variant | 23.4 | 25.98 | 16.5 | 25.95 | 20.87 | 11.97 | 0.38 |
| splice_acceptor_variant | 0.32 | 0.23 | 0.26 | 0.31 | 0.2 | 0.15 | 0 |
| splice_donor_variant | 0.33 | 0.36 | 0.27 | 0.43 | 0.11 | 0.14 | 0.01 |
| splice_region_variant | 1.18 | 1.22 | 1.04 | 0.94 | 1.16 | 1 | 0.06 |
| start_lost | 0.03 | 0.03 | 0.02 | 0.03 | 0.01 | 0.02 | 0 |
| stop_gained | 1.21 | 1.96 | 0.88 | 2.03 | 2.71 | 0.79 | 0.02 |
| stop_lost | 0.02 | 0.01 | 0.01 | 0.01 | 0.03 | 0.01 | 0 |
| stop_retained_variant | 0.01 | 0.02 | 0.01 | 0.01 | 0.01 | 0.01 | 0 |
| synonymous_variant | 10.74 | 10.56 | 8.96 | 11.9 | 5.92 | 7.02 | 0.22 |
| upstream_gene_variant | 13.26 | 11.8 | 14.14 | 13.13 | 10.83 | 13.47 | 9.64 |

### Gene-motif differential analysis in each subtype

Our gene association analysis revealed that there are several genes that are significantly mutated in multiple subtypes. This observation encouraged us to find out if mutations in these genes have different motif preferences in each subtype. To do this, we investigated the mutations in tri-nucleotide motifs in the significant genes for each subtype. We found a considerable number of genes were significantly mutated in multiple subtypes, and that the mutations happened in different tri-nucleotide motifs. An example of this phenomena is *TP53* which was identified as a significant gene in both SC1 and SC5. However, the gene was identified to have different gene-motif patterns in each of these subtypes. The preference tri-nucleotide motif of mutations in SC1 is CT-G.G, however, the preference tri-nucleotide motif of mutations in SC5 is CT-T.G. *KRAS* is another example where most of its mutations occurred in CT-A.C and CT-G.C for SC1; CA-A.C and CT-A.C for SC2; CT-A.C, CA-A.C, CT-G.C for SC3; CT-G.C, CA-A.C, CT-A.C for SC4; CT-G.C, CT-G.T, and TG-A.T for SC5; CT-A.C, CT-G.C, CA-A.C for SC6; and CT-A.C, CA-A.C, CA-C.A for SC7 (**Figure 4a)**. A full list of genes that significantly mutated in multiple subtypes but in different tri-nucleotide motifs, is provided in **Supplementary table S6.**

**Figure 4.**
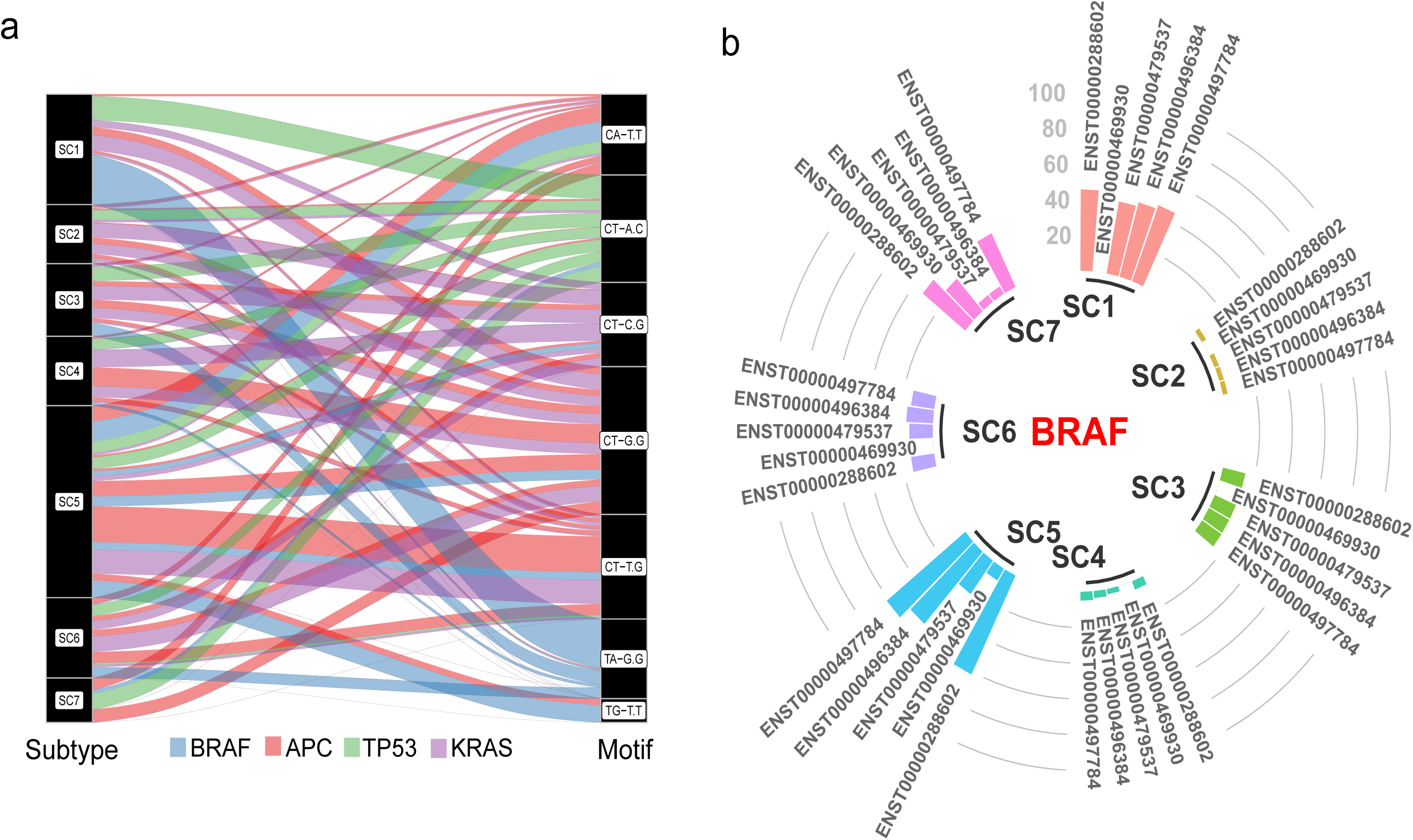
Enrichment of mutations in different motifs and transcripts. **a)** Investigating the mutations in tri-nucleotide motifs of genes that become significant in multiple subtypes revealed a motif context specificity of mutations in each subtype. For example, *TP53* becomes a significant gene in both SC1 and SC5. However, the preference tri-nucleotide motif of mutations in SC1 is CT-G.G, but in SC5 is CT-T.G. **b)** Our analysis also showed that for some of the candidate genes, the mutations are enriched in a specific transcript of the gene.

### Mutation rate in transcripts

We then investigated the difference between our identified subtypes with respect to the mutational load in different transcripts of the coding genes. Our analyses revealed that for many of the candidate coding genes, the mutations occurred in specific transcripts of the genes (**Figure 4b** and **Supplementary figure S12**). Interestingly, 68.11% of coding genes were significantly mutated in multiple subtypes, however, the mutations enriched in different transcripts of the genes (**Supplementary table S7**). For example, gene *TTN* is significantly mutated in all CRC subtypes except SC7. Interestingly, 62% of samples in SC5 mutated in transcript *ENST00000436599* which is much higher than other subtypes (7%, 1%, 6%, 5% and 12% for subtypes SC1, SC2, SC3, SC4, and SC6 respectively). In addition, for gene *PCDHA2* that is significantly mutated in subtypes SC1, SC5 and SC7 we observe different patterns. The *De Novo* mutations for this gene in SC5 are enriched in ENST00000378132 and ENST00000520672, while in SC1 and SC7 the mutations are enriched in transcript ENST00000526136 (**Supplementary figure S12).** We provide a list of candidate genes with enrichment of *De Novo* mutations in different transcript of the gene in **Supplementary table S7.**

### Gene ontology and pathway analyses

Next we conducted gene ontology (GO) and gene pathway analyses to investigate whether candidate genes in the identified subtypes are associated with any specific gene ontology or pathway term [46-48]. To do this, we chose 50 highly significant genes in each subtype and performed GO and pathway analyses using WebGestalt [49] (*see method section*) for each subtype. As shown in **Figure 5a** several unique and common GO terms have been identified for each subtype. For example, “cellular response to interferon-gamma (GO:0071346)” was only associated with SC3, “cell development (GO:0048468)” and “regulation of cell motility (GO:2000145)” unique to SC7, and “self-proteolysis (GO:0097264)” uniquely associated with SC5. Conversely, “cell adhesion (GO:0007155)” is an example of common gene ontologies enriched in subtypes SC1, SC4 and SC5, and “regulation of postsynaptic membrane potential (GO:0060078)” common in SC2, SC5 and SC7. A full list of ontology terms that significantly associated with each subtype is provided in **Supplementary table S8.** Pathway analysis also identified 30 pathways that are associated with subtypes SC3, SC4 and SC6 (**Figure 5b**). Similar to GO terms, we also identified several unique pathways associated with different subtypes. For instance, SC4 has a unique pathway called “Signaling by *FGFR3* point mutants in cancer” that has been previously associated with urothelial, breast, endometrial, squamous lung cancers and ovarian cancer [50]. Additionally, “Interferon gamma signaling” are represented in SC3 and SC6, which can be used in immunotherapy for CRC [51]. For more details, look at the **Supplementary table S9.**

**Figure 5.**
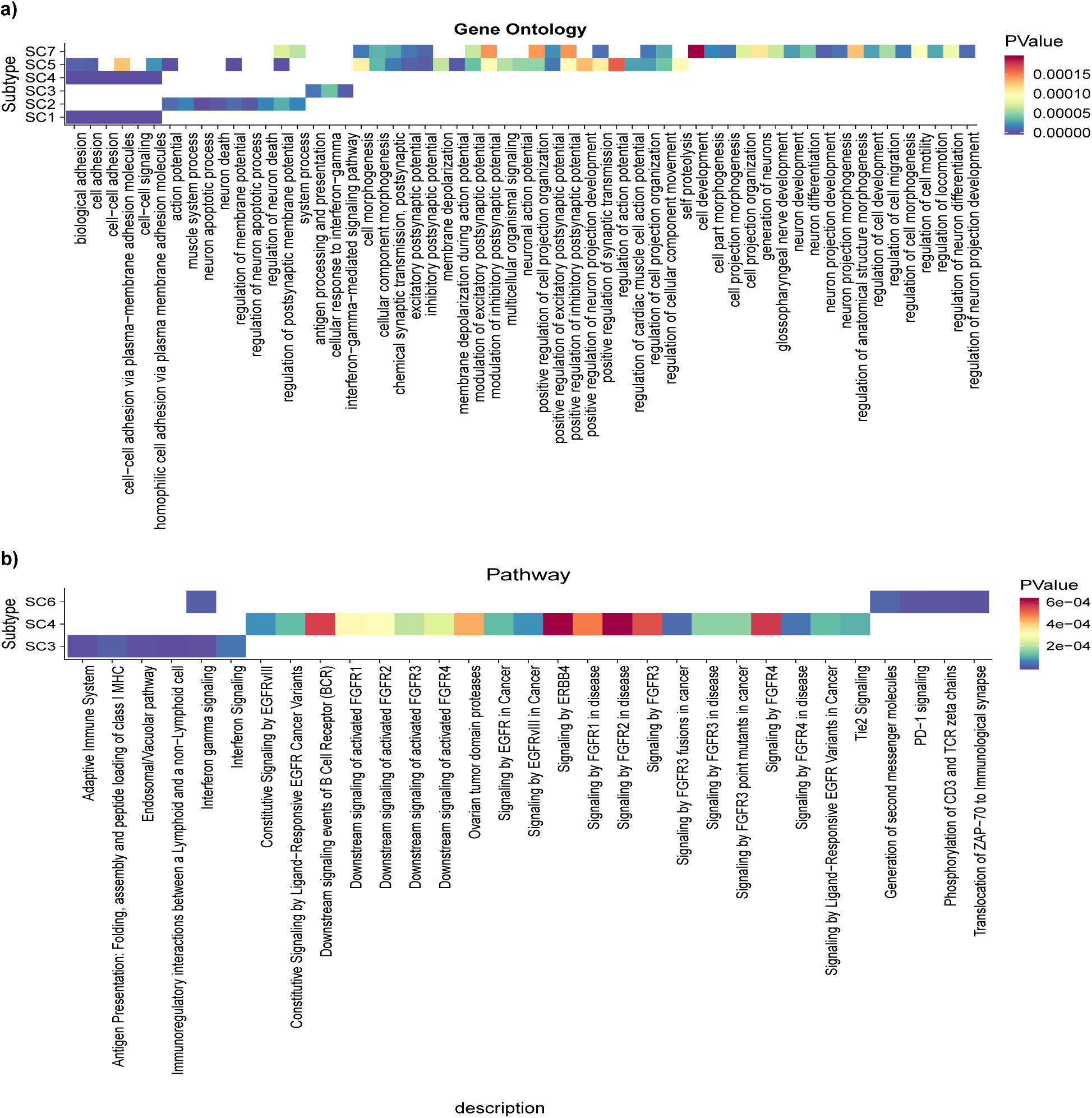
Gene ontology and pathway analysis. **a)** Gene Ontology analysis. Applying the WebGestalt tool [49] on 50 highly mutated coding genes in each subtype revealed several unique and common gene ontologies in the CRC subtypes. For example, “cellular response to interferon-gamma (GO:0071346)” is only associated to SC3. Also “Cell development (GO:0048468)” and “regulation of cell motility (GO:2000145)” ontologies are uniquely associated to SC7. A full list of ontology terms that were significantly associated with each subtype is provided in **Supplementary table S9. b)** Gene pathway analysis. Our gene pathway analysis also revealed 30 pathways that were associated with subtypes SC3, SC4 and SC6. Again, we identified several unique pathways associated to different subtypes. For example, SC4 has a unique pathway called “Signaling by FGFR3 point mutants in cancer” that has been previously associated with urothelial, breast, endometrial, squamous lung cancers and ovarian cancer [50]. For more details about gene pathway analysis look at **Supplementary table S9.**

### Clinical report and survival analysis

We also examined clinical data, such as gender, ethnicity, and regions that were available for a subset of the CRC samples. Our findings demonstrate interesting patterns in our data allocated in each subtype, such as the fact that 52.3% of the patients in SC7 are female and that 76.9% and 60.9% of patients are male in SC5 and SC3, respectively (see more detail in **Figure 6a** and **Supplementary table S10**). Geographical differences were also observed in each subtype, such as all the 21 patients in SC7 being from China and 68.2% of patients in SC1 are from US (e.g. COAD project; **Figure 6b**). Additionally, our analysis also showed that samples in SC5 are older than those in SC2. This observation is shown by the histogram plot for each subtype based on age in **Supplementary figure S13**. We also found that most of the samples in SC4 had cancer, at the proximal and distal of the colon, especially in the rectum and cecum and ascending colon sites.

**Figure 6.**
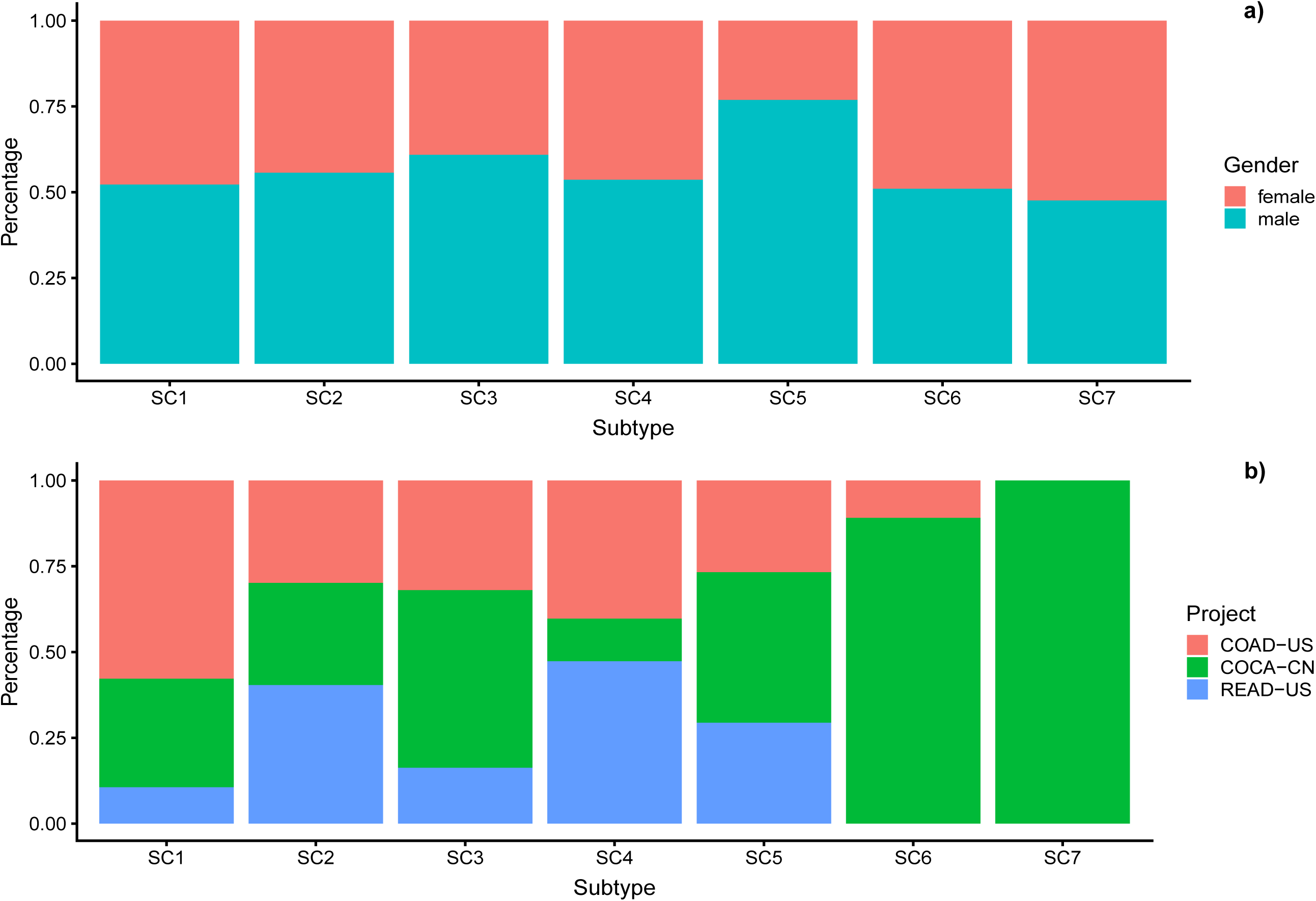
Gender and geographical distributions of the patients in each subtype. **a) Gender distributions**. Our gender analysis showed that 52.3% of the patients in SC7 are female, 76.9% and 60.9% of patients in SC5 and SC3 are male, respectively. **b) Geographical distributions**. Our geographical distribution analysis showed that 21 patients in SC7 are from China and 68.2% of patients in SC1 are from US (e.g. COAD project).

Considering the molecular data for a subset of CRC samples, we also performed a survival analysis. In many cases, we had the data of how many days patients were alive after joining this study for sample collection. We created survival curves for all subtypes using the Kaplan-Meier method [52] and compared the survival curves using log-rank test and recorded those with P-value less than 0.0001. Our survival analysis illustrates that SC3 patients have more chance to survive and SC7 has a short survival duration compared to the other subtypes. The Kaplan-Meier plot for survival and the P-values of a log rank test has is shown in **Figure 7**.

**Figure 7.**
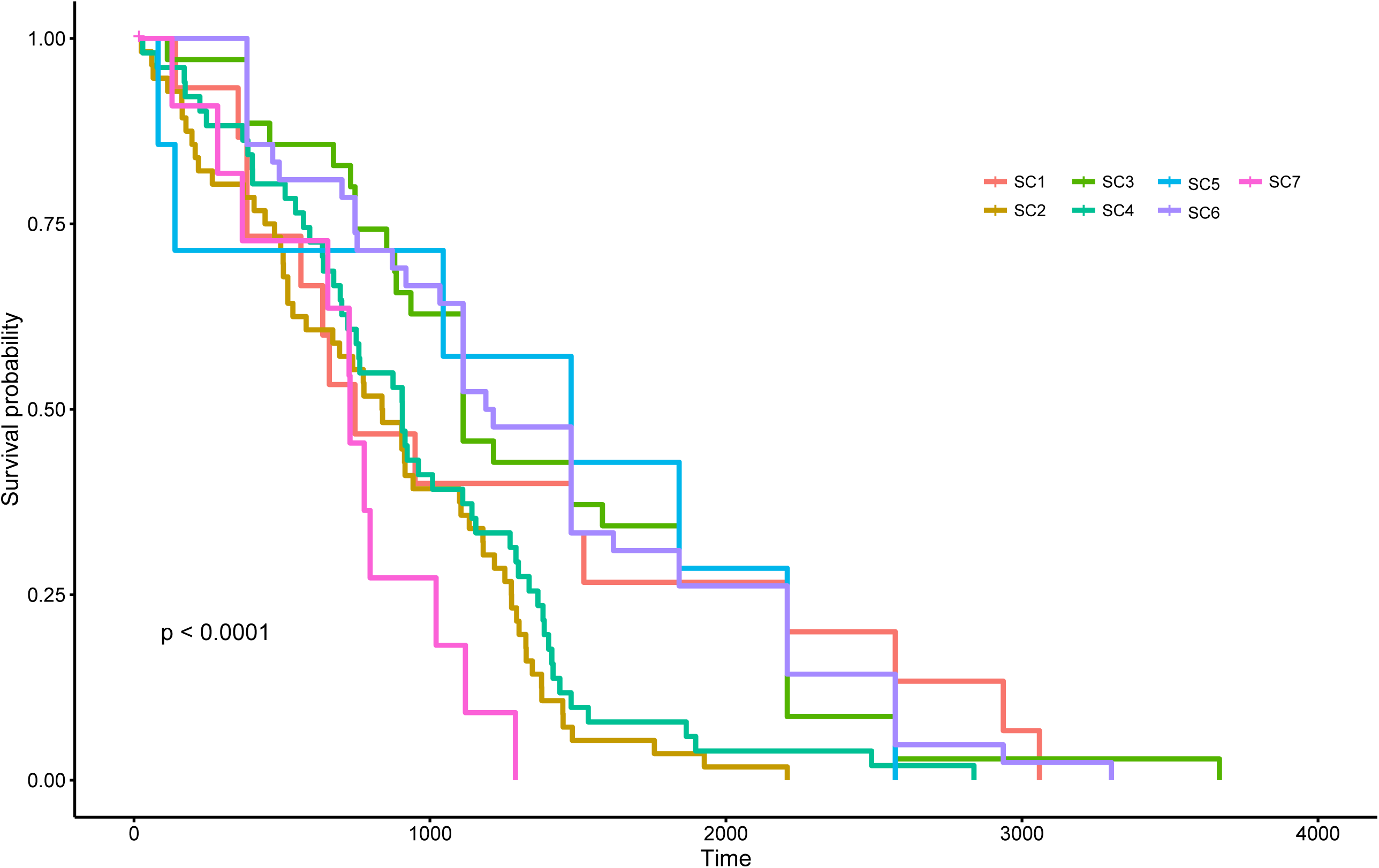
Survival analysis. Using the Kaplan-Meier method we compared survival curves of the CRC subtypes. The analysis illustrated that SC3 patients have more chance to survive and SC7 have short survival duration compared to the other subtypes.

## Discussion

Classification of cancers into subtypes is essential because it informs personalized medicine therapeutic and/or prevention strategies. The CRC Subtyping Consortium introduced four consensus molecular subtypes (CMS1–4) for CRC, based on gene expression profiles of tumors. Recently, a study by Lochlan *et al*. [53] investigated genome-scale DNA methylation in a large cohort of CRC patients and revealed five distinct CRC subtypes of colorectal adenocarcinomas with different therapeutic biomarkers for CRC treatment. This highlights the need for a better and more inclusive subtype classification approach to enable biomarker discovery and thus inform drug development for CRC.

Mutation is the hallmark of cancer genome, and many studies have reported cancer subtyping based on the type of frequently mutated driver genes [8, 9, 54], or the proportion of mutational processes [9], however, none of these existing cancer subtyping methods consider these features simultaneously. In other words, the sequence context of somatic point mutations in driver genes have not been taken into consideration in cancer subtyping and biomarker discovery. Here, we integrated these two features (frequently mutated genes and sequence context of mutated sites) and implemented a bioinformatics pipeline for CRC subtyping using the ICGC whole genome and whole exome somatic point mutations. Our analyses revealed 3,131 driver gene-motifs, which were mutated at a significantly higher level in CRC samples compared to other cancer samples. Using 3,131 gene-motifs as features, we identified seven unique CRC subtypes, which are associated with distinct symptoms and genotypes in CRC. As **Figure 2** shows some of the gene-motifs are subtype specific. For example, we found that 43% of SC7 samples had ACG-to-ATG mutations in *HDAC9*, while other subtypes had no mutation in this motif within *HDAC9*. 40% of SC5 samples had TCG-to-TTG mutations within *HDAC9*, but no other subtypes had a significantly elevated mutation in this gene-motif (i.e. maximum rate of mutation for other subtypes in this gene-motif was 5%). Another example is *BRAF* for which we found that 41% of samples in SC1 had GTG-to-GAG mutations. This is significantly higher than other CRC subtypes (1%, 11%, 2%, 0% 8%, and 0% for SC2 to SC7, respectively). Interestingly, none of the samples from SC5, which is a hypermutated subtype, had mutations in *BRAF* (**Supplementary table S3**). Importantly, in some cases, most of the subtypes are significantly mutated in a gene, but these mutations are enriched in different trinucleotide contexts. For example, Grb2-associated binder 1 (*GAB1*) is a docking protein, which is strongly implicated in CRC. This gene is significantly mutated in SC1, SC3, SC5, SC6 and SC7. However, the mutations are enriched in CCG-to-CTG for SC1 (31%), ACT-to-ATT for SC3 (100%), TCT-to-TAT for SC5 (35%), TTG-to-TCG for SC6 (67%), and GCA-to-GTA for SC7 (100%). These data suggest that, the sequence context of mutations within driver genes (frequently mutated genes) contains invaluable information that can be used to accurately identify CRC subtypes.

Our investigation of coding and lncRNA genes also revealed that there is a different mutational load for coding and lncRNA genes in CRC subtypes (Supplementary figures S8 and S9). Our analysis indicated that 74% of cancer mutations fall within lncRNAs, yet their role in driving tumorigenesis is not well understood. Our data showed that mutational load in some lncRNAs can be as high as protein coding genes (**Supplementary figure S10**). Specifically, for the CRC subtype SC7, which includes tumors with less mutated genomes, the average rate of mutations for coding and lncRNA genes are very similar (**Supplementary figure S10**). Similar to protein coding genes, somatic mutations in highly mutated lncRNAs are also sequence context specific. For example, we show that the well-known CRC-associated lncRNA *TTN-AS1* is highly mutated in all subtypes, except SC7. We found that mutations CTG->CGG, CTG->CAG, and CAT->CTT are enriched in SC1, SC3, and SC5, respectively (**Supplementary table S6**).

Additionally, our analyses revealed that for 68.11% of candidate coding genes, somatic mutations were enriched in different transcripts of the genes (**Figure 3b; Supplementary figure S12**, and **Supplementary table S7**). To our knowledge, this result represents the first account of transcript-specific mutations in cancer and may provide a new insight into cancer driver mechanisms.

We found that, on average, all CRC subtypes had small levels of mutational signatures 1 and 3 (**Figure 3a**), both of which reported for CRC patients [44]. Our data indicate that only a small subset of tumors within each subtype show highly increased levels of these two types of mutations. It has been suggested that the mutational process underlying these signatures (a combination of these signatures represents COSMIC signature 10) is altered activity of the error-prone polymerase POLE [44]. We also found that mutational signature 2 in our study (**Figure 3b**) is a common among all CRC subtypes. This signature is reported to be associated with an endogenous mutational process derived by spontaneous deamination of 5-methylcytosine. In contrast to Signatures 1,2, and 3, some of the other signatures showed significantly different levels in different CRC subtypes. For example, Signature 5 showed greater exposures in SC4 and SC5. There is no known etiology for this signature. Signature 6 exhibited greater exposures in SC3 and SC7. This signature is likely associated with defective DNA mismatch repair [44]. Altogether, some but not all the differences present in tumors in terms of the extent of mutational processes are reflected in our subtypes.

Our survival analyses show that subtype 7 (SC7) is associated with a poor outcome (**Figure 7**). There are 100 gene-motifs (from 13 different genes) and at least 50% of samples in SC7 showed significantly elevated mutations in these gene-motifs, while other subtypes had no mutation (or showed very mutations with low frequencies) in these gene-motifs. Interestingly, most of these 13 genes (*KMT2C, DPP6, PRIM2, SDK1, PTPRN2, CNTNAP2, CSMD1, CTNNA3, MAGI2, EYS, HLA-DRB1, DLGAP2, FANK1*) are known to be associated with cancer. For example, somatic mutation in Cub and Sushi Multiple Domains 1 (*CSMD1*) gene is associated with an early age of presentation in CRC individuals [55]; or mutation in *PTPRN2* (Receptor-type tyrosine-protein phosphatase N2) can promote metastatic and cellular migration in breast cancer cells through lipid-dependent sequestration of an actin-remodeling factor [56]. Our analysis also showed that SC2 and SC4 had very similar survival curves. Importantly, the gene-motif profiles of these two subtypes are also very similar (**Figure 2**). For example, all samples in both SC1 and SC4 subtypes had mutation in the processed transcript *XXbac-BPG254F23*. Interestingly, none of samples in other subtypes had mutation in *XXbac-BPG254F23* (**Supplementary table S6**).

## Conclusions

In conclusion, with genomic medicine emerging as a routine part of the healthcare system and cancer individuals now frequently sequenced for specific sets of mutations, a large amount of whole genome and whole exome sequencing samples have become available for analysis. Accurate classification of cancer individuals with similar mutational profiles may help clinicians to accurately identify individuals who could receive the same types of treatment. Here, by application of a unique statistical pipeline and a novel “gene-motif” concept, and integration of somatic mutations from the ICGC consortium, we have identified seven subtypes with strong biological characterization in CRC. Looking to the future, screening for these subtypes has the potential to provide an accurate identification of the cancer-associated genes. By knowing the genes and phenotypes associated with the subtypes, a personalized treatment plan can be developed that considers the specific phenotypes associated with their genomic lesion. Our list of subtype-specific genes provides a better understanding of the underlying genetic causes of CRC. More importantly, for the first time we provided a system-wide analysis of the enrichment of *denovo* mutations in a specific motif context of the genes in CRC. By knowing the genes and motif associated with the mutations, a personalized treatment plan can be developed that considers the specific motif context of mutations within responsible genes. Lastly, the pipeline developed in this study is freely available and will be useful in the analysis of cancers and in narrowing down the causative genes within each cancer.

## Method and material

### Overall design

The flowchart of our pipeline is shown in **Figure 1**. At the first step, mutations are annotated with the FANTOMCAT robust gene list. In the next step, we find a distribution representing somatic mutations data and compute the P-values for each gene and gene-motif based on negative-binomial and beta-binomial distributions, respectively and find significant genes and gene-motifs using a standard threshold of P-value< 0.01. Then, we use model-based clustering to identify subtypes in the CRC samples based on significant gene-motifs. Finally, we identify biomarkers and interpret biological characterization in each subtype.

### ICGC Dataset

We used the ICGC dataset, which contains data from 19 types of cancers including CRC. It contains 536 patients with CRC across three projects (READ-US, COAD-US, COCA-CN) from China and US where 45% are female and 55% are male. 96% of these data were captured by whole genome sequencing (WGS) technology and the rest of them by non-NGS technology.

### FANTOMCAT coding and non-coding genes list

We used a robust gene list of the FANTOMCAT consortium which contains 21,069 protein-coding genes and 27,919 lncRNAs. FANTOMCAT used cap analysis of gene expression (CAGE) technology to conduct high-confidence coding and non-coding genes. These genes are sturdily approved by transcription initiation evidences. Therefore, we decided to annotate mutations based on this dataset instead of other consortiums.

### Statistical pipeline to identify significant genes

We identified coding and non-coding genes that significantly mutated in the colorectal samples in the following manner. We first counted the number of samples that had mutation in each gene. We next used a negative-binomial distribution as the best fit distribution for our data (**Supplementary figure S2**) to identify significant genes. Finally, we calculated the P-value for each gene using the following formula: 

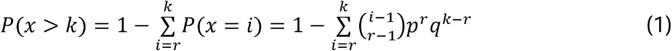

Using the mentioned pipeline, we identified 382 coding genes (listed in the **Supplementary table S1**) that significantly mutated in the colorectal samples. We also identified beta-binomial distribution as the best fit compared to other methods to identify gene-motifs that significantly mutated in the colorectal samples. The **Supplementary figure S3** shows the fit diagrams of different distributions on the gene-motifs. We then computed the P-values for each gene-motif as follow: 

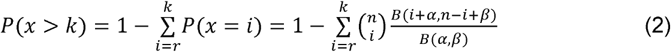

where *B*(*α,β*) is the beta function. Using the mentioned pipeline, we identified 3,131 significantly mutated motifs within 382 significant coding genes (listed in **Supplementary table S2**). We called this concept “gene-motif”. We used the R package “VGAM” [57] to perform the above analyses.

### Clustering

We used model-based clustering to identify subtypes in colorectal samples. Model-based clustering is one of the density-based unsupervised machine learning methods which is non-parametric. Hence, it does not consider any assumption for the number of available clusters. This feature is suitable here, as the number of subtypes of CRC is unknown and it is better to do clustering without any assumptions. From a statistical perspective, each sample is independent with respect to its number of mutations. Hence, via the central limit theorem we expect that if subtypes are correctly identified, the mutational load in determined features come from a Gaussian distribution. Model-based clustering automatically fits the best Gaussian distribution to samples, and finds the optimal number of clusters [15]. Here, we use model-based clustering in a hierarchical manner. Using the clustering method, we identified two clusters with 450 and 83 patients at the first clustering level. These two clusters in turn are divided into smaller and still distinguishable subgroups in the second clustering level.

### Biological characterization

#### Mutational signature analysis

We used CANCERSIGN [58] to identify signatures that are represented in our CRC samples. CANCERSIGN uses a non-negative matrix factorization (NMF) algorithm to identify optimal number of signatures. As it was shown in **Supplementary figure S14**, in overall we identified seven signatures in CRC samples. The exposure matrix for identified signatures are also provided in **Supplementary table S11**.

#### Gene, lncRNA, and transcript rates analysis

We used Fisher’s exact test to identify coding and lncRNA genes that significantly mutated in each subtype. To do this, we made a contingency table as follow: number of CRC samples with mutation in the gene; number of CRC samples that had no mutation in the gene; number of samples with other cancer with mutation in the gene; number of samples with other cancers that had no mutation in the gene. The results are shown in **Supplementary file S1**. We did the same process for transcripts to identify transcripts that significantly mutated in each subtype.

#### Consequence type of mutations

The consequence type of mutations from the ICGC dataset have been used to count the consequence type of each mutation. We then calculated the relative frequency of consequence types in each subtype.

#### Gene ontology analysis and gene pathway analysis on the significantly mutated coding genes

We used the gene ontology analysis tool WebGestalt [59] to observe the over-representation of gene ontology and pathways associated with significant genes in each subtype, separately. We used default values for WebGestalt parameters (FDR < 0.05, 2000 permutation, minimum gene = 5). We also used the top 50 genes associated with each subtype as the candidate genes.

#### Clinical information

We downloaded clinical data for the CRC samples from ICGC (http://cancer.digitalslidearchive.net). Two metadata files “sample.tsv” and “donor.tsv” have been used for analysis of gender and age of patients.

#### Survival analysis

We used the Kaplan-Meier method to conduct survival curves for all subtypes. We used “survival” [60] and “survminer” [61] R packages to conduct Kaplan-Meier curves and obtain the significance of survival prediction for subtypes. Long-rank test was also applied to obtain the P-value for survival analysis.

## Author contributions

HAR designed the study. HD, HAR, and AD developed hypothesis and experiments (the problem definition, candidate gene-motif identification, clustering, and hierarchical model. HAR, HD, and AD wrote the manuscript; HD, HAR, AD, HRR, DE, JB, NL edited the manuscript; HD carried out the analyses including the statistical analyses, candidate gene and gene-motif identification, text mining, gene prioritization, gene ontology, and survival analysis, HRR helped with the statistical analyses, MB performed mutational signatures analysis. HD generated all figures and tables. All authors have read and approved the final version of the paper.

## Data availability

The source code has been attached to this paper as **Supplementary file 2**, the somatic point mutations can be downloaded from ICGC website.

## Acknowledgements

HAR is supported by a UNSW Scientia Program Fellowship. HAR was previously a research associate in Harry Perkins Institute of Medical Research under supervision of Prof Alistair Forrest. Analysis was made possible with computational resources provided by the Telethon Kids Bioinformatics Server with funding from the Australian Government and the Government of Western Australia. HRR is supported by IRN National Science Foundation (INSF) Grant No. 96006077.

## Supplementary tables legends

**Table S1.** List of 382 significantly mutated genes that were identified by negative-binomial distribution.

**Table S2.** List of 3,131 significantly mutated gene-motifs that were identified by beta-binomial distribution.

**Table S3.** List of CRC patients that were grouped in each subtype using a model-based clustering.

**Table S4.** Frequency of mutation in 3,131 significantly mutated gene-motifs in each subtype.

**Table S5.** Subtype-specific genes that were identified by a Fisher exact test. Rare mutated genes in each subtype also highlight by green color.

**Table S6.** List of protein-coding genes that become a significant gene in different subtypes but with enrichment of mutations in different motif context.

**Table S7.** Fraction of mutations in different transcripts of candidate genes in each subtype.

**Table S8.** Gene ontology analysis from WebGestalt tool [49] on 50 highly mutated coding genes in each subtype. We only considered those ontology terms with FDR <= 0.05.

**Table S9.** Gene pathway analysis from WebGestalt tool on 50 highly mutated coding genes in each subtype. We only considered those terms with FDR <= 0.05.

**Table S10.** Gender analysis on CRC samples in each subtype.

**Table S11.** The exposure matrix for identified signatures in our CRC samples.

## Supplementary figures legends

**Figure S1.**
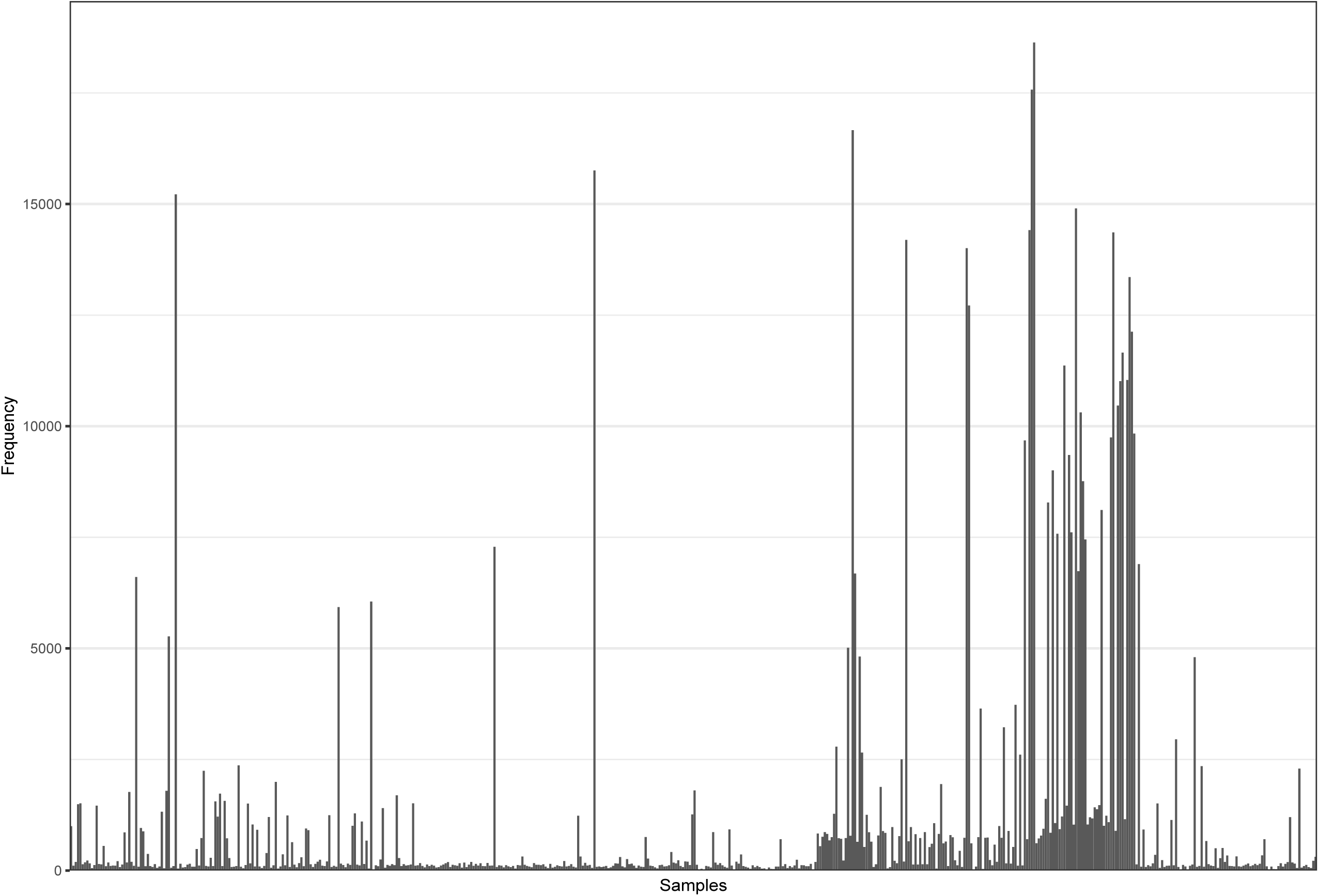
Mutation rates of the CRC patients. This plot shows number of mutations in each CRC sample.

**Figure S2.**
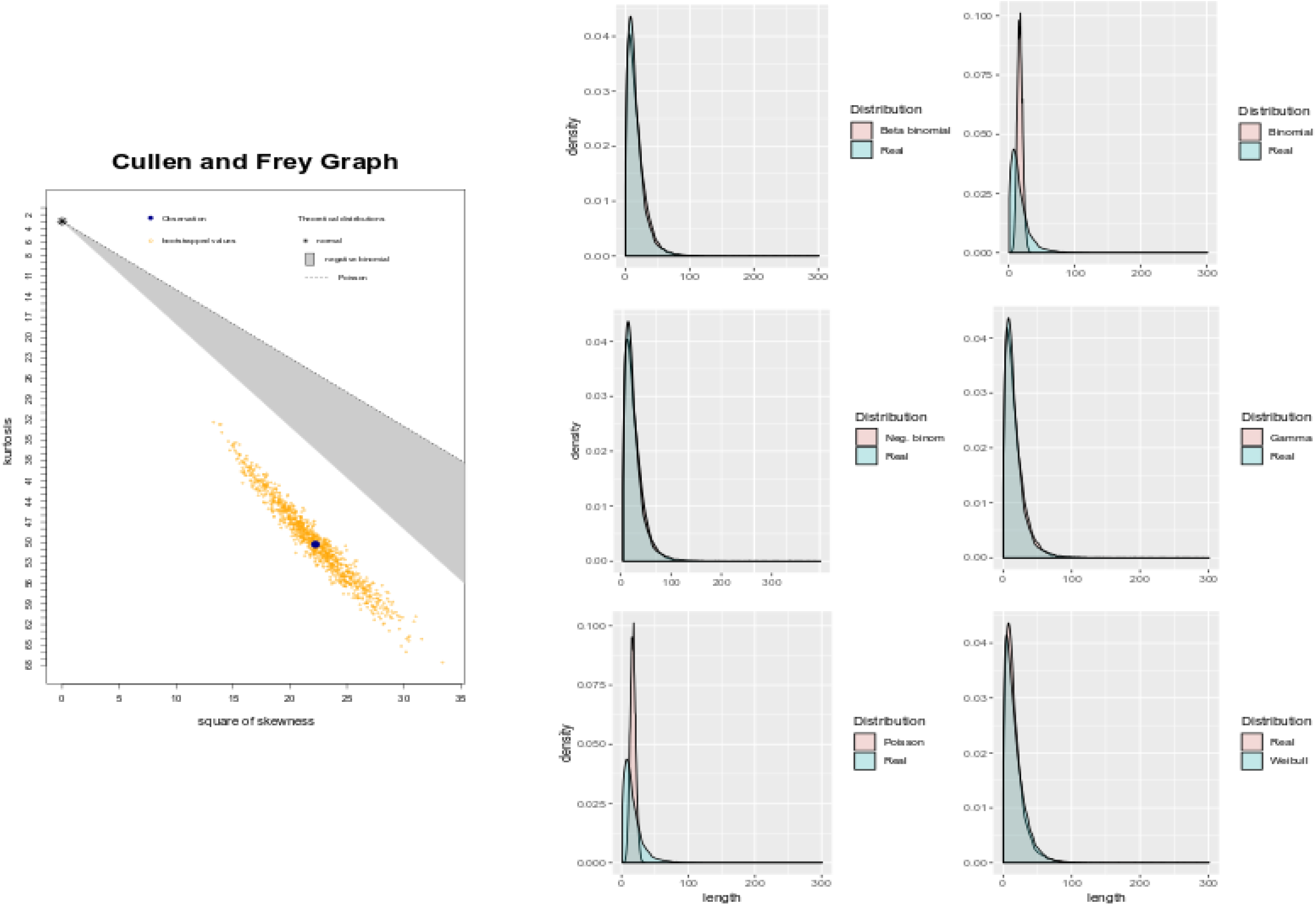
Identify best-fitted distribution to discover significant genes. This plot shows our comparison of different distribution techniques to fit the number of mutations in the genes and identify significantly mutated genes.

**Figure S3.**
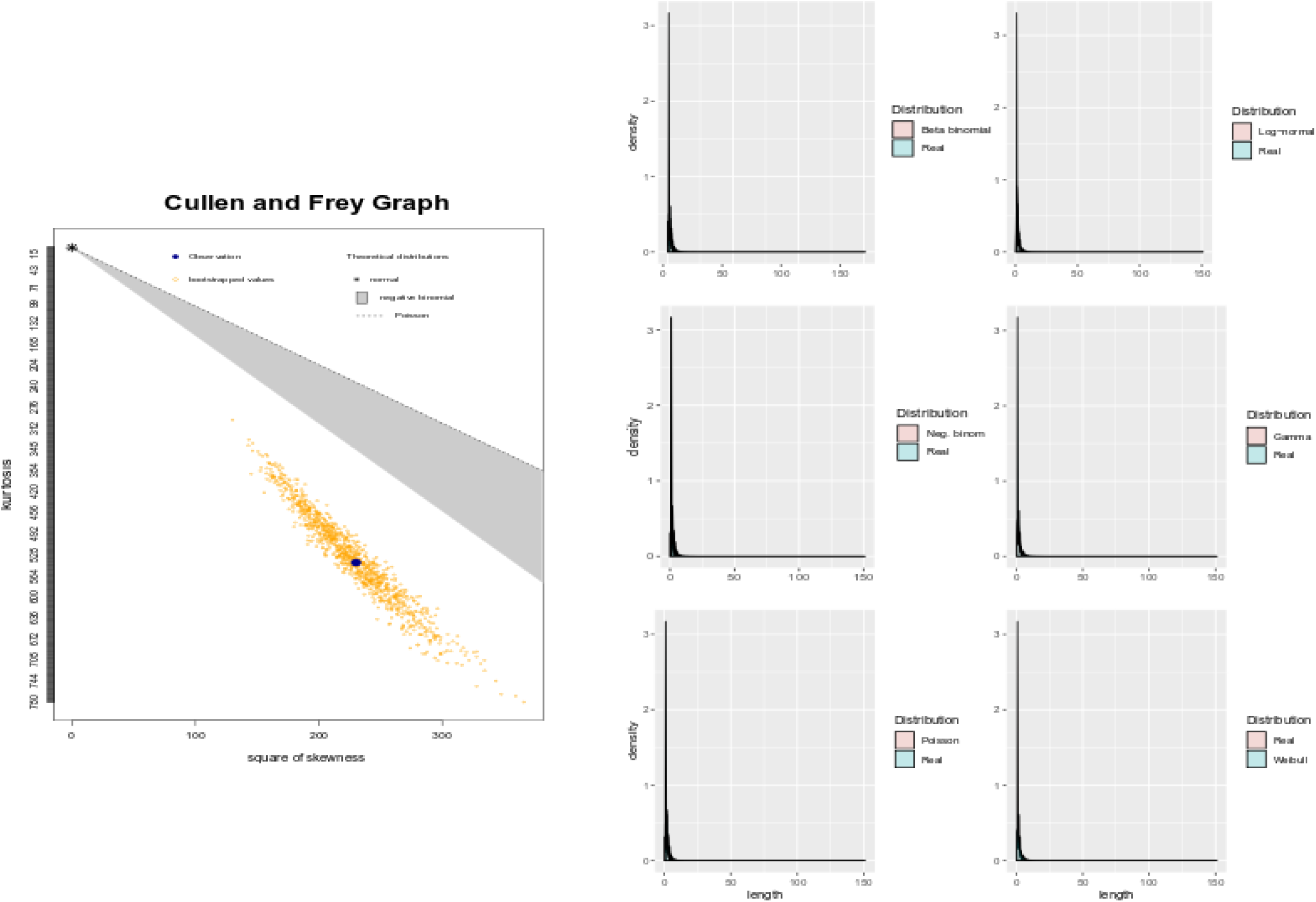
Identify best-fitted distribution to discover significant gene-motifs. This plot shows our comparison of different distribution techniques to fit the number of mutations in the gene-motifs and identify significantly mutated gene-motifs.

**Figure S4.**
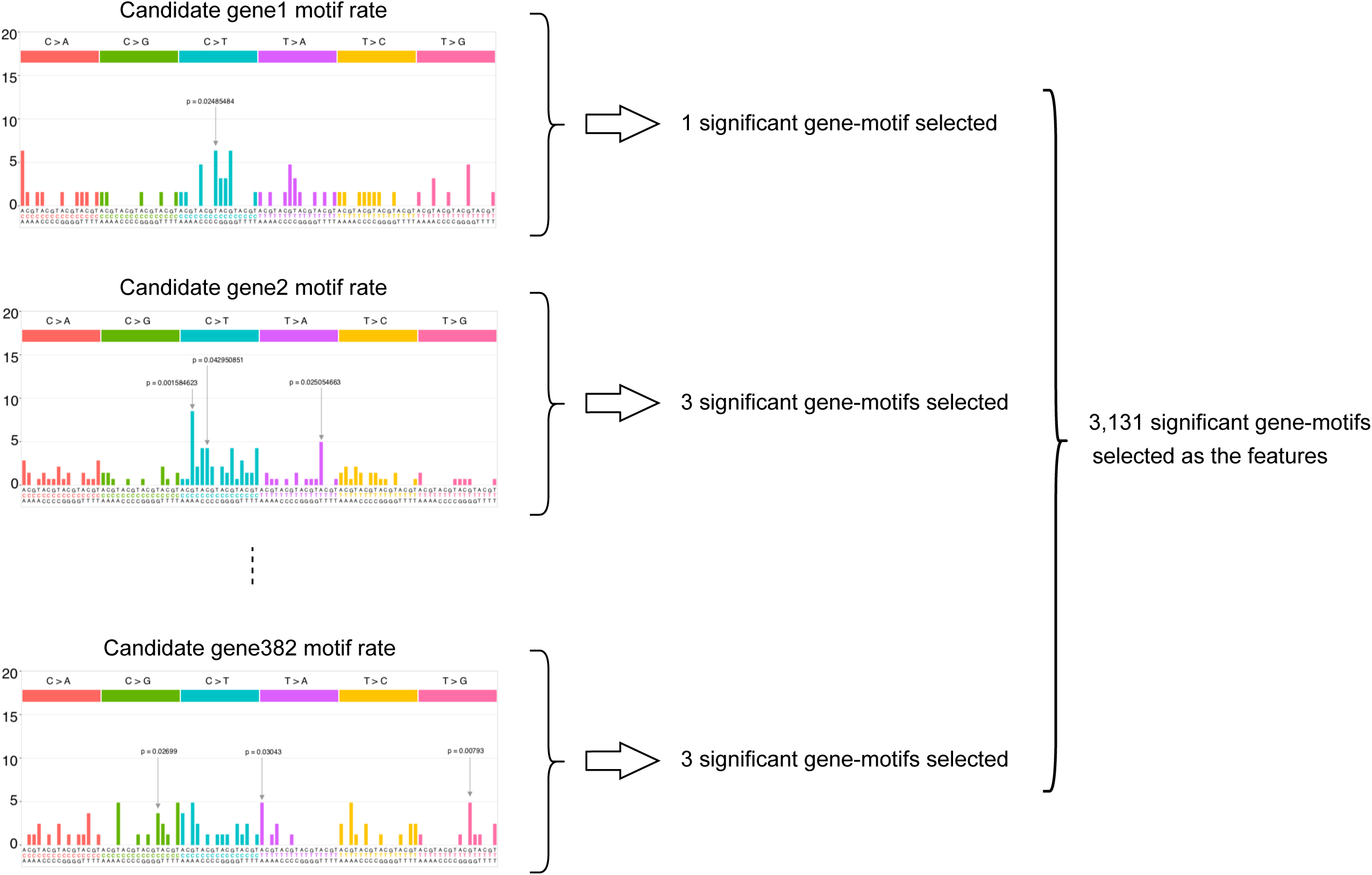
An overview of gene-motifs concept. We first identify 382 significantly mutated coding genes in colorectal cancers (candidate genes). We then used Fisher exact test to identify those motifs that significantly mutated within candidate genes.

**Figure S5.**
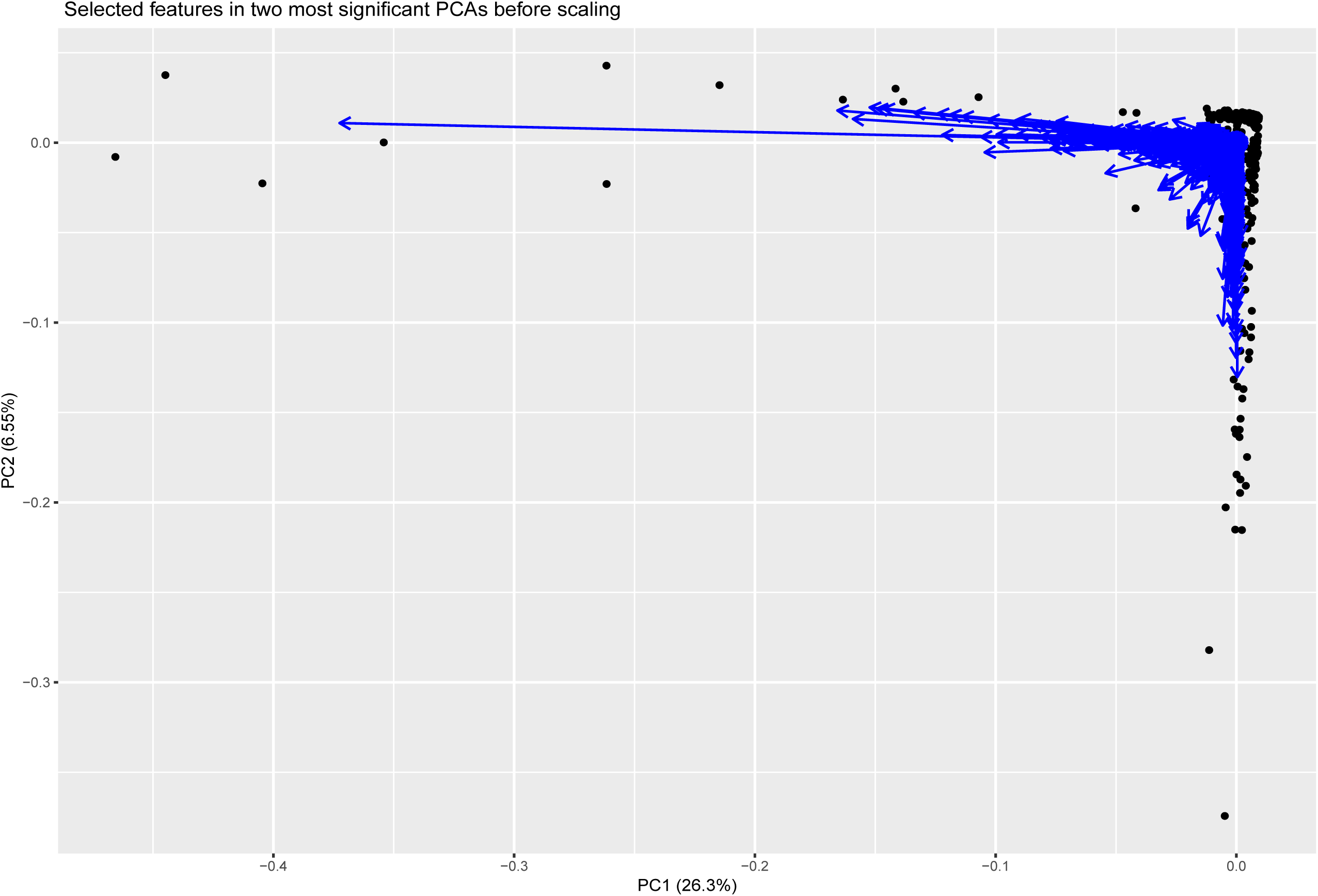
Selected 3,131 features in two most significant PCAs before scaling. This plot shows two principal components (PCs) that demonstrates the potential discrimination that can be obtained from our identified features.

**Figure S6.**
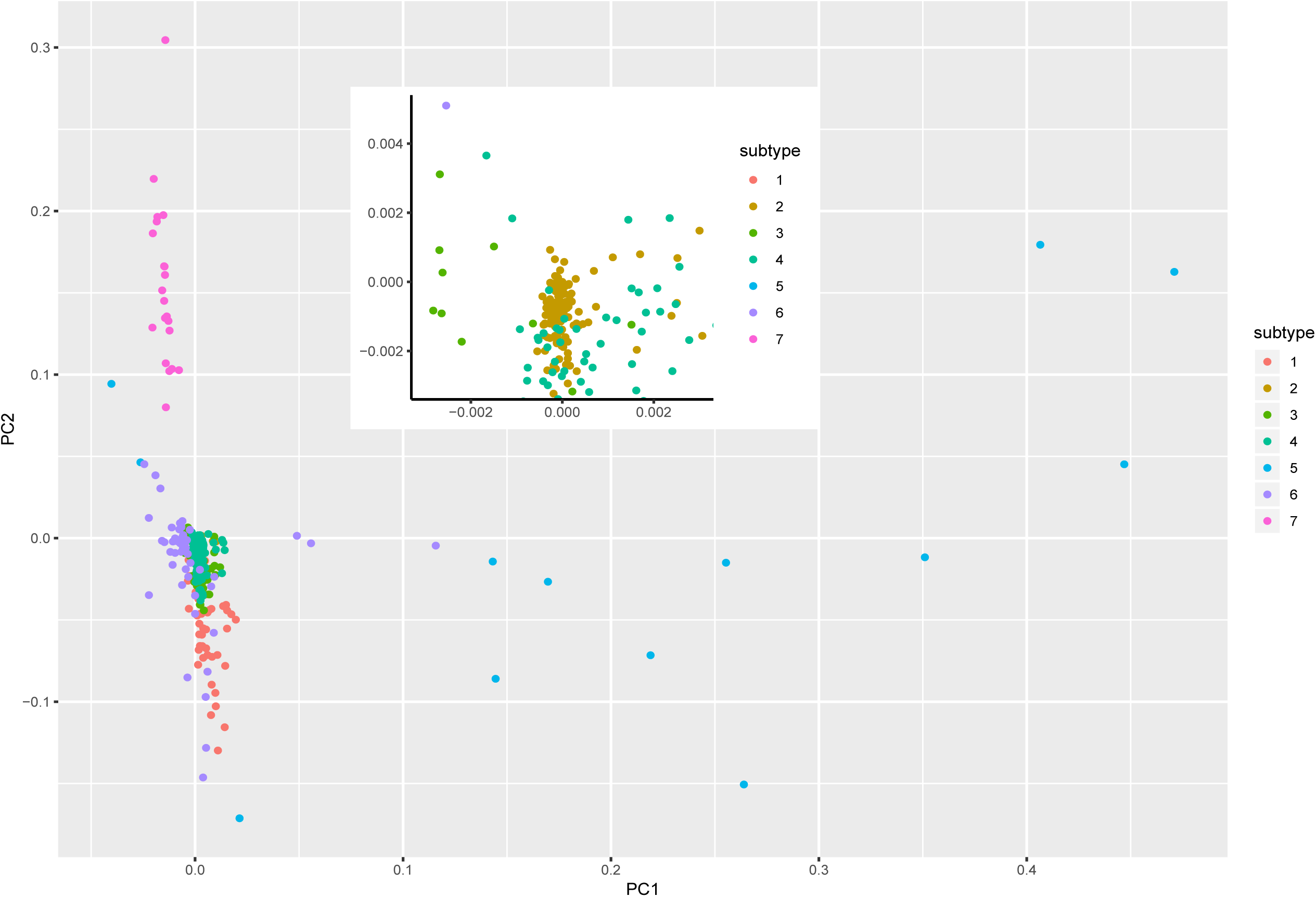
Illustration of patients in two first PCAs of features. Distribution of CRC samples through 3,131 gene-motif features by PCA analysis.

**Figure S7.**
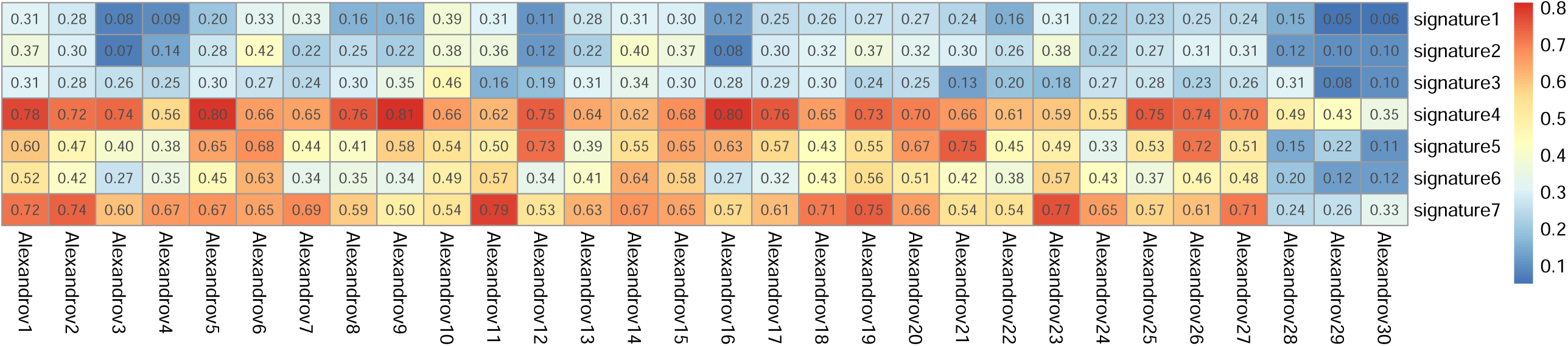
Correlation between our identified signatures and Alexandrov’s signatures in each CRC subtype separately.

**Figure S8.**
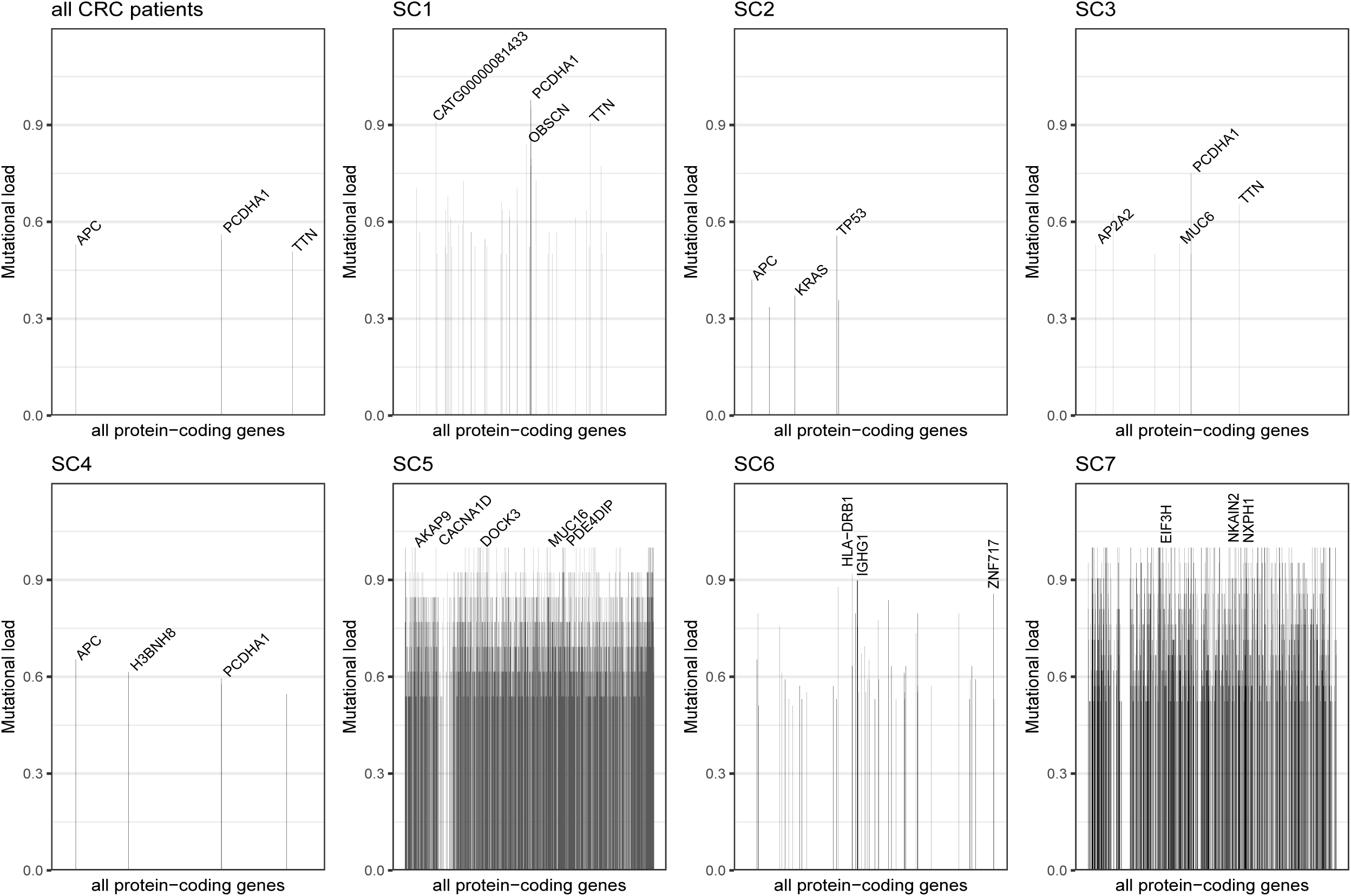
Mutational load of protein coding genes in each subtype separately. Each bar chart shows fraction of samples with mutation in a gene.

**Figure S9.**
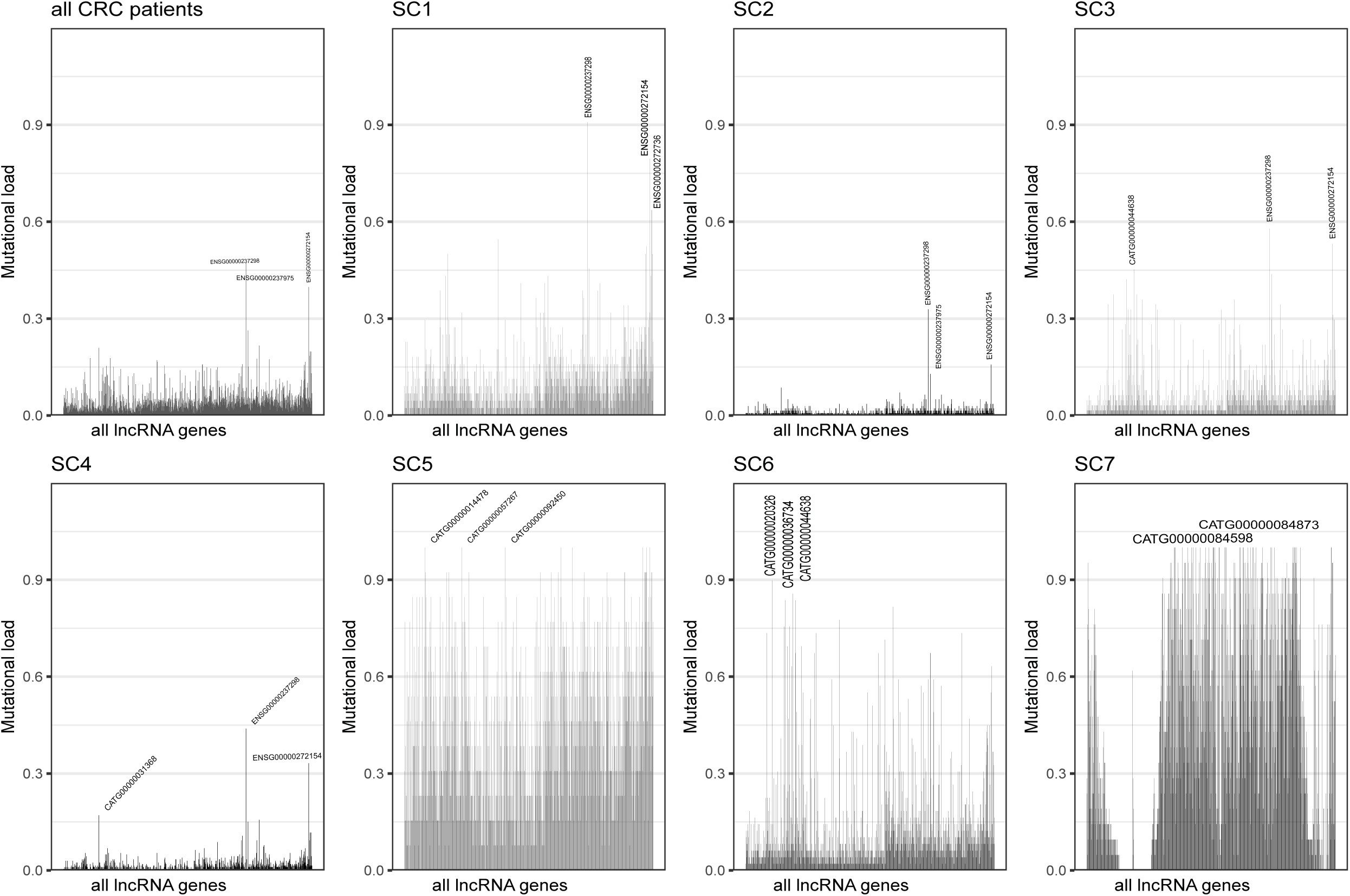
Mutational load of long non-coding RNA genes in each subtype separately. Each bar chart shows fraction of samples with mutation in a lncRNA.

**Figure S10.**
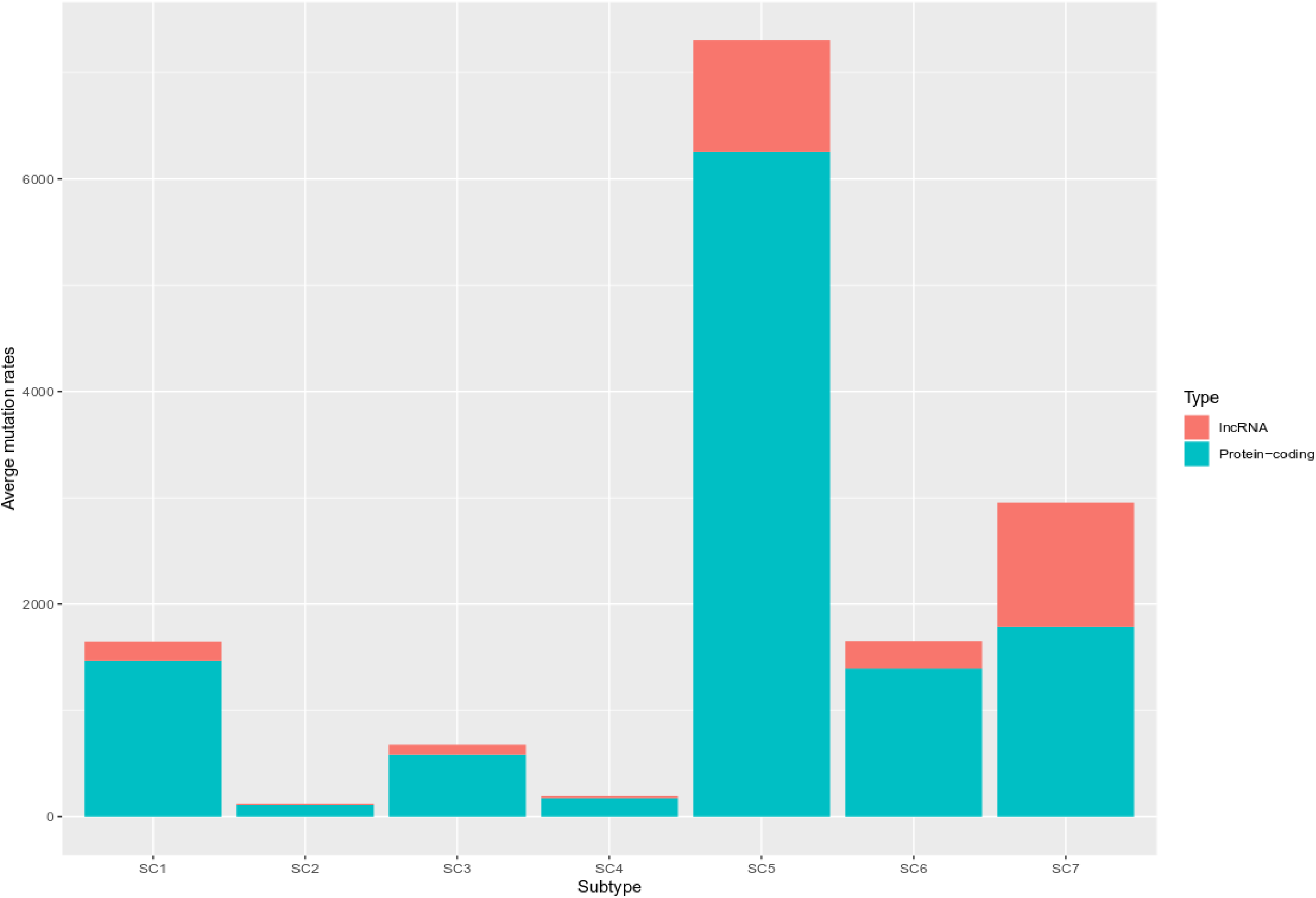
Mutation rates in coding and lncRNA genes in each subtype. Red color indicates average number of mutations in lncRNA genes and green color indicates average number of mutations in coding genes.

**Figure S11.**
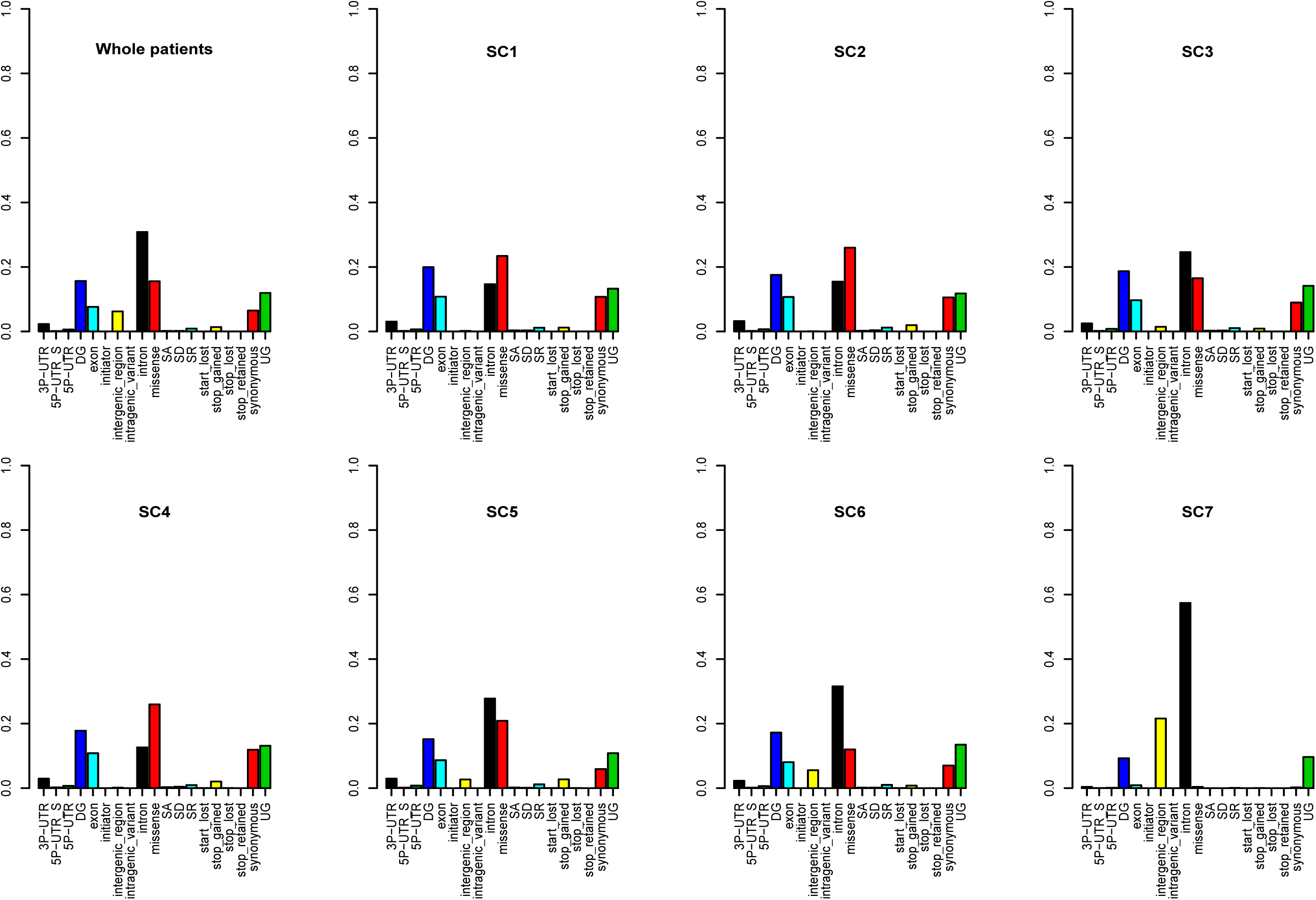
Consequence type analysis. Figure shows fraction of mutations in different consequence types for each subtype.

**Figure S12.**
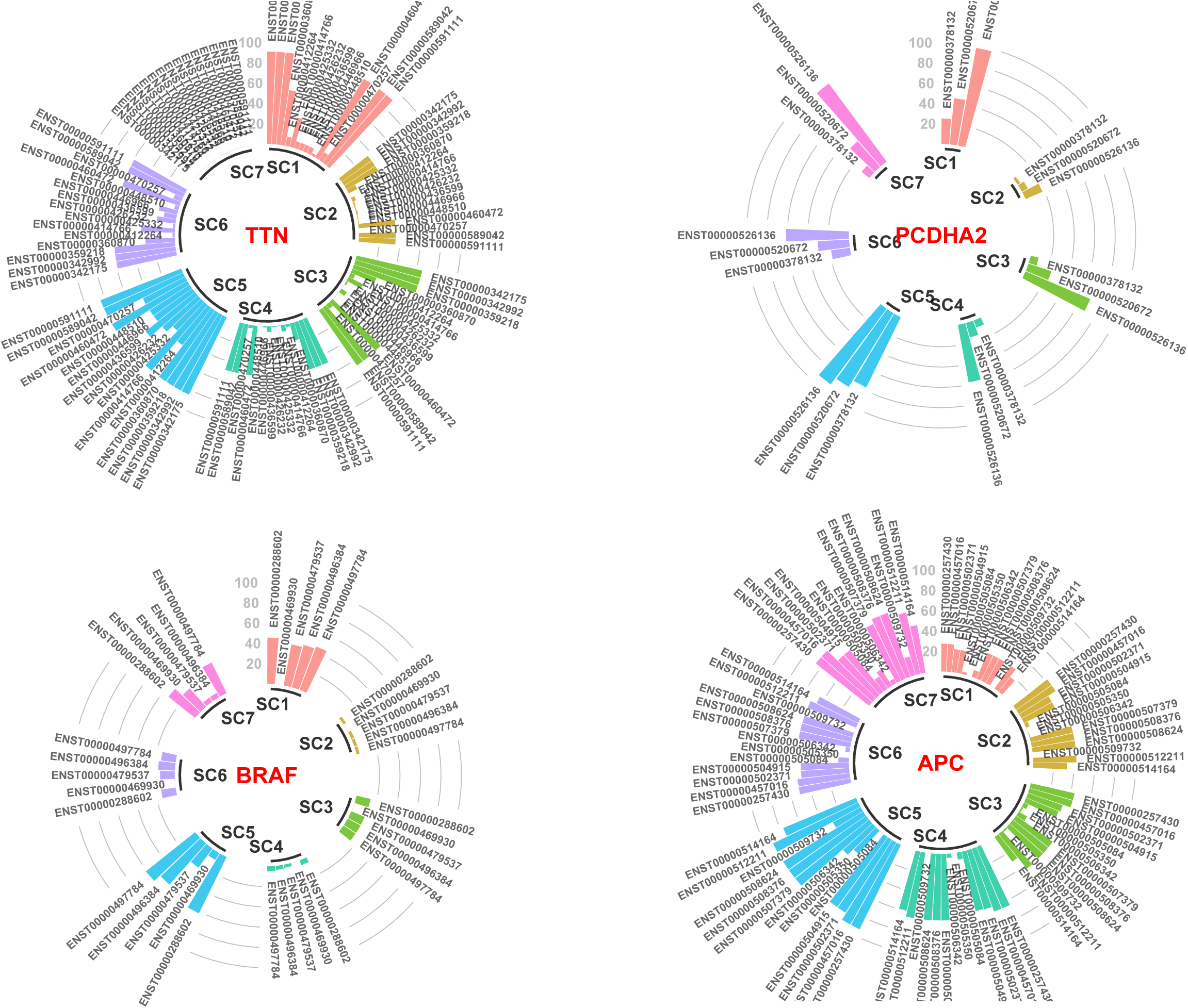
Mutation rate in transcripts in genes TTN, PCDHA2, BRAF, APC. Figure shows mutational rate in different transcripts of genes *TTN, PCDHA2, BRAF, and APC* across the CRC subtypes identified in this study.

**Figure S13.**
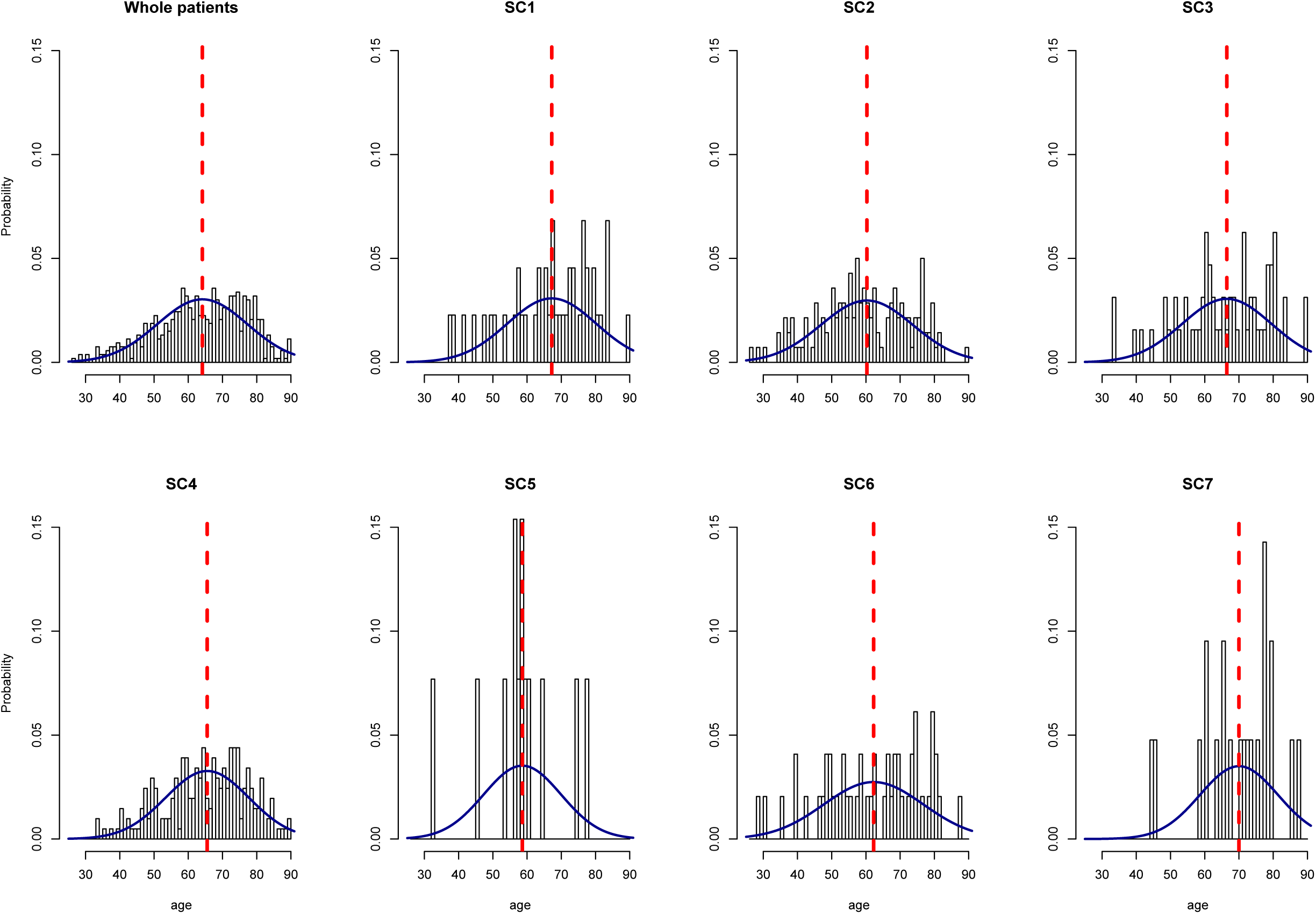
Analysis of age distribution of CRC samples in the identified subtypes.

**Figure S14.**
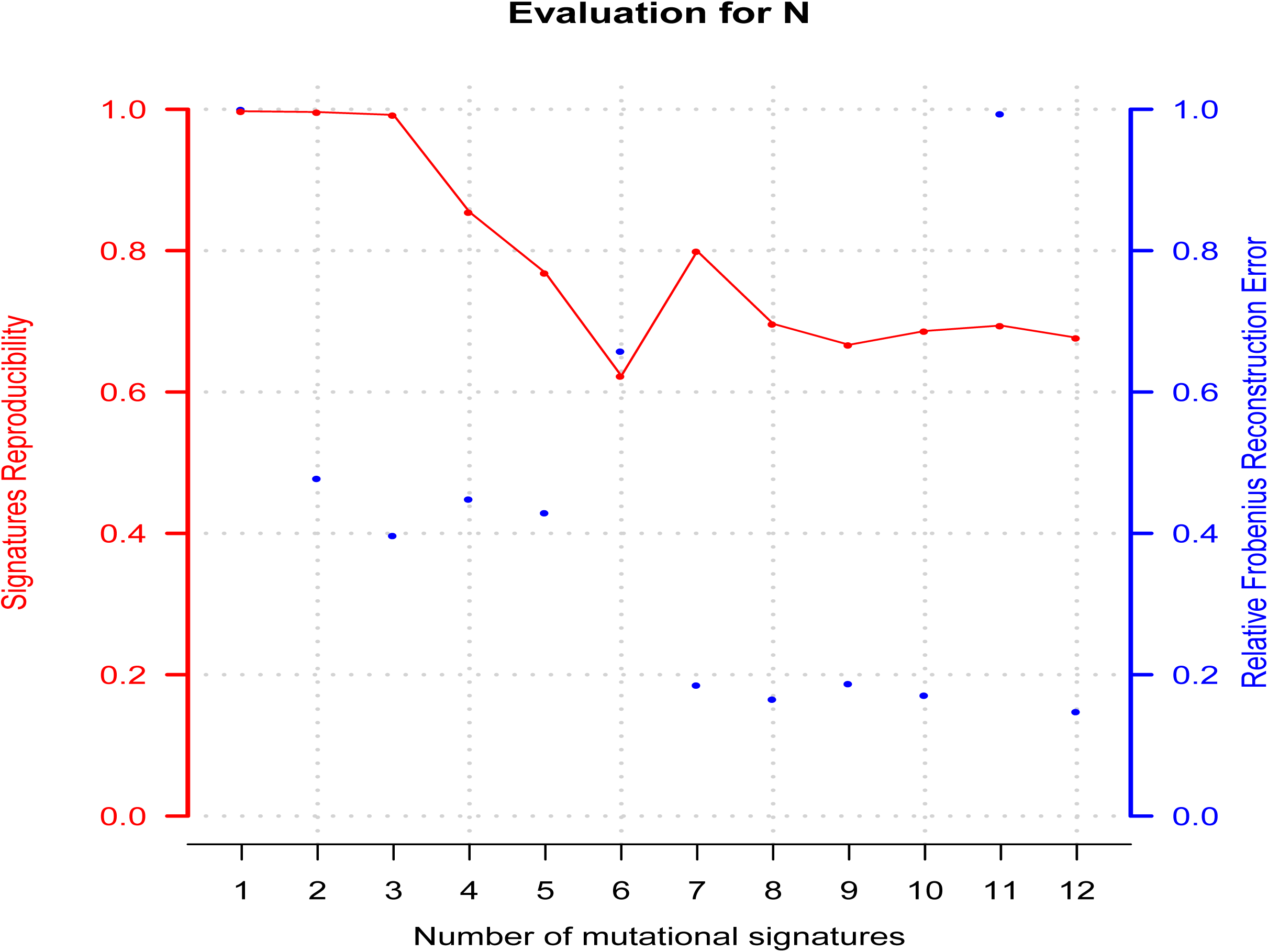
Evaluation plot for deciphering 3-mer mutational signatures in the CRC samples. We used the CANCERSIGN tool [45] to identify mutational signatures in CRC samples. The evaluation plot of deciphering 3-mer mutational signatures become optimized for seven signatures.

